# DSSNA: Automated Nucleic Acid Structure Analysis in Molecular Dynamics

**DOI:** 10.64898/2026.09.25.754511

**Authors:** S. V. Gorelov, A. L. Konevega, A. V. Shvetsov

## Abstract

This work presents DSSNA (Define Spatial Structure of Nucleic Acids) v2026, a free and open-source standalone GROMACS[1] module for the automated analysis of nucleic acid structures from atomic coordinates that reproduces the core functionality of X3DNA[2] and DSSR[3] while extending it to the analysis of molecular dynamics (MD) trajectories. The algorithm identifies and classifies canonical and non-canonical base pairs, hydrogen-bond networks, stacking interactions, helices, and other elements of nucleic acid secondary and tertiary structure, using neighbor-search algorithms for efficient analysis of large trajectory datasets and a modular output system that lets users select the structural metrics required for a particular analysis. Application to a 1 *µ*s MD trajectory of a GUAA tetraloop mutant of the sarcin–ricin domain of *Escherichia coli* 23S ribosomal RNA (PDB: 1MSY) showed that persistent canonical base pairs form the structural core that maintains the secondary-structure profile, whereas most non-canonical base pairs and stacking interactions are transient. These results demonstrate that DSSNA enables the characterization of both static nucleic acid structures and the temporal evolution of structural interactions in MD simulations.

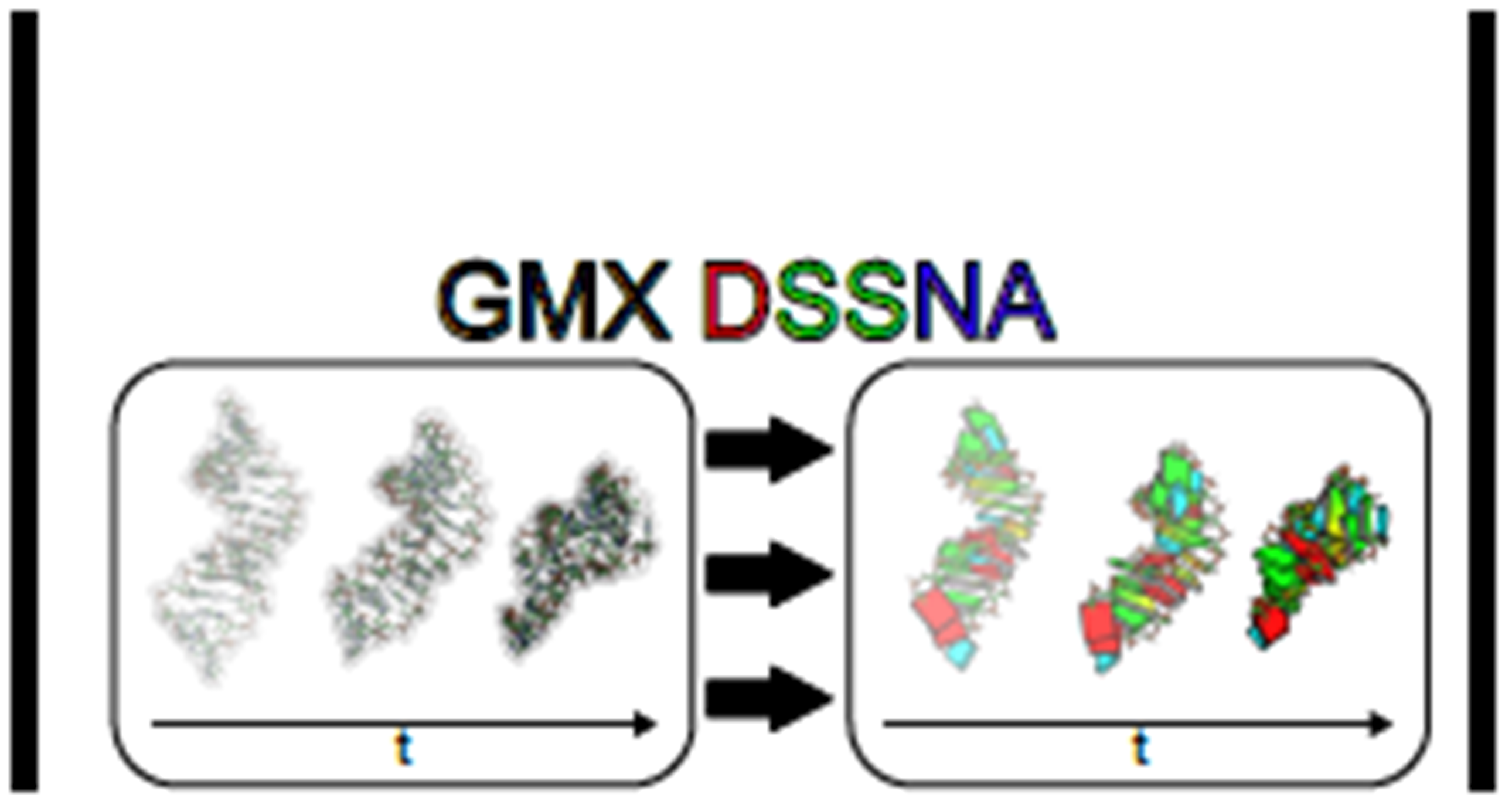

## Introduction

Structural analysis of nucleic acids is a key area of molecular biology and bioinformatics, as the three-dimensional organization of DNA and RNA directly determines their functional properties. Accurate algorithms for describing secondary and tertiary structures not only enable the interpretation of experimental data but also facilitate in-depth analysis of dynamic processes, including folding, interactions with proteins, and ligand-mediated conformational transitions.

Contemporary nucleic acid research extends far beyond the description of static structures. Increasing attention is being paid to the dynamics of conformational states, with the aim of elucidating the mechanisms underlying the regulation of genetic processes, the recognition of RNA motifs, and the structural basis of mutagenesis. Addressing these questions requires tools capable of automatically and reproducibly analyzing the temporal evolution of spatial interactions across large molecular dynamics datasets.

The development of such algorithms is of fundamental importance for addressing a wide range of challenges, from modeling mutations that affect RNA and DNA stability to identifying novel drug targets and designing nucleic acid nanostructures. The development of accurate, openly available analytical methods will contribute to the establishment of a reproducible computational infrastructure for structural biology and make the results of molecular simulations accessible to the broader scientific community.

Currently available tools for nucleic acid analysis, such as X3DNA and DSSR, provide a comprehensive set of functions for identifying and classifying structural elements of nucleic acids. However, their application in contemporary computational studies—particularly in the analysis of molecular dynamics trajectories—remains limited. The main challenges include restricted access to source code, difficulties integrating these tools with widely used analysis packages, and substantial computational costs associated with processing large datasets. These limitations highlight the need for open, flexible, and computationally efficient tools for the automated analysis of nucleic acid structures in dynamic systems.

In this study, we present DSSNA (Define Spatial Structure of Nucleic Acids) v2026, a standalone GRO-MACS module developed to determine secondary and tertiary structure from atomic coordinates, including the analysis of time series generated by molecular dynamics simulations. DSSNA was strongly inspired by the functionality and conceptual framework of X3DNA/DSSR, but it was developed as an independent implementation rather than as a direct port of their source code. DSSR is distributed as a licensed closed-source software package, whereas X3DNA provides source code for some of its releases through a registration-based distribution system and under project-specific licensing conditions. Consequently, the internal implementation details and source-code organization of DSSR were not available for direct inspection. The development of DSSNA therefore relied on published scientific articles, technical documentation, user manuals, tutorials, and the publicly observable behavior of X3DNA/DSSR. This approach enabled us to reproduce the core structural-analysis concepts of these tools while developing an independent implementation integrated with GROMACS. Therefore, DSSNA was developed as an openly available implementation with the aim of improving transparency, extensibility, and integration with GROMACS-based molecular dynamics workflows. This makes DSSNA a useful tool for researchers working with both individual structures and long-timescale molecular dynamics simulations.

## Implementation

For the algorithm to operate correctly, the user must provide two types of input files: a molecular structure or molecular dynamics trajectory, and a binary topology file generated during the preprocessing stage. The algorithm then prompts the user to select the atom group containing the nucleic acid to be analyzed (this step can be skipped if the user specifies the desired group with the *−sel* flag). Before analyzing the frames of an MD trajectory — or a static structure — DSSNA validates the input parameters provided by the user. Before iterating over the frames of a molecular dynamics trajectory, it is necessary to determine which molecule will be analyzed, including, at a minimum, its molecular composition. Since the molecular dynamics force fields and simulation frameworks used in contemporary software packages generally do not account for the formation or breaking of covalent bonds, it is reasonable to assume that the chemical structure of the molecule remains unchanged over time. Therefore, the developed algorithm parses the topology before analyzing any trajectory frame in order to determine the composition of the molecule. More specifically, it identifies the nucleotides that constitute the molecule.

Nucleotides in DSSNA are identified by analyzing the presence of key base-ring atoms characteristic of purine and pyrimidine residues. This approach is robust against modifications in the base, sugar, or phosphate moieties, as it relies primarily on the geometry and atom names of the base ring. The nucleotide-type assignment is performed only once, during the analysis of the topology prior to the analysis of trajectory frames. Modified nucleotides are automatically mapped to their closest canonical counterparts for the purpose of structural classification, while their original residue names from the topology are retained in the output. This mapping strategy is analogous to the approach used in DSSR[3], where modified nucleotides are assigned one-letter abbreviations corresponding to their canonical bases while retaining information about their modified status in the output. The mapping of modified residues to standard bases ensures compatibility with established structural-analysis conventions and facilitates the interpretation of results.

The algorithm processes the frames of a molecular dynamics trajectory individually and analyzes each frame independently of the others. Once the analysis is complete, it writes the results to the output files specified by the user. These files contain the information selected by the user — for example, helices, base pairs, hydrogen-bond networks, or other structural features — for the selected region of the trajectory. By default, the entire trajectory (or static structure) is analyzed.

The determination of nucleic acid secondary structure requires the rapid and efficient identification of hydrogen bonds (see S.V) and base-stacking interactions (see S.XII), which form the basis for detecting more complex structural patterns. This, in turn, requires structural objects located within a certain distance of one another to be identified efficiently. To address this type of problem, DSSNA uses the Neighbor Search algorithm implemented in GROMACS[1].

Rather than producing an unnecessarily large amount of information whenever a particular metric is reported, we designed the output system to be modular. The only required standard output is information about nucleotide frames (see S.I). To obtain more detailed information — such as the geometric characteristics of a base pair or the hydrogen-bond network between the corresponding nucleotides — the user must explicitly request an output option that enables these metrics.

## Example usage

As an application example, the developed algorithm was used to analyze a 1 *µ*s MD trajectory of a GUAA tetraloop mutant of the sarcin–ricin domain of *Escherichia coli* 23S ribosomal RNA (PDB: 1MSY), demonstrating how trajectories can be analyzed using a subset of the features provided by DSSNA (see Figure 1). Since 1MSY was used as a reference structure in the DSSR user manual[3], we selected it to demonstrate the results obtained by applying the developed DSSNA algorithm to the MD trajectory of this system. Details of the MD simulation protocol are provided in the Molecular Dynamics section of the Supporting Information.

**Figure 1.**
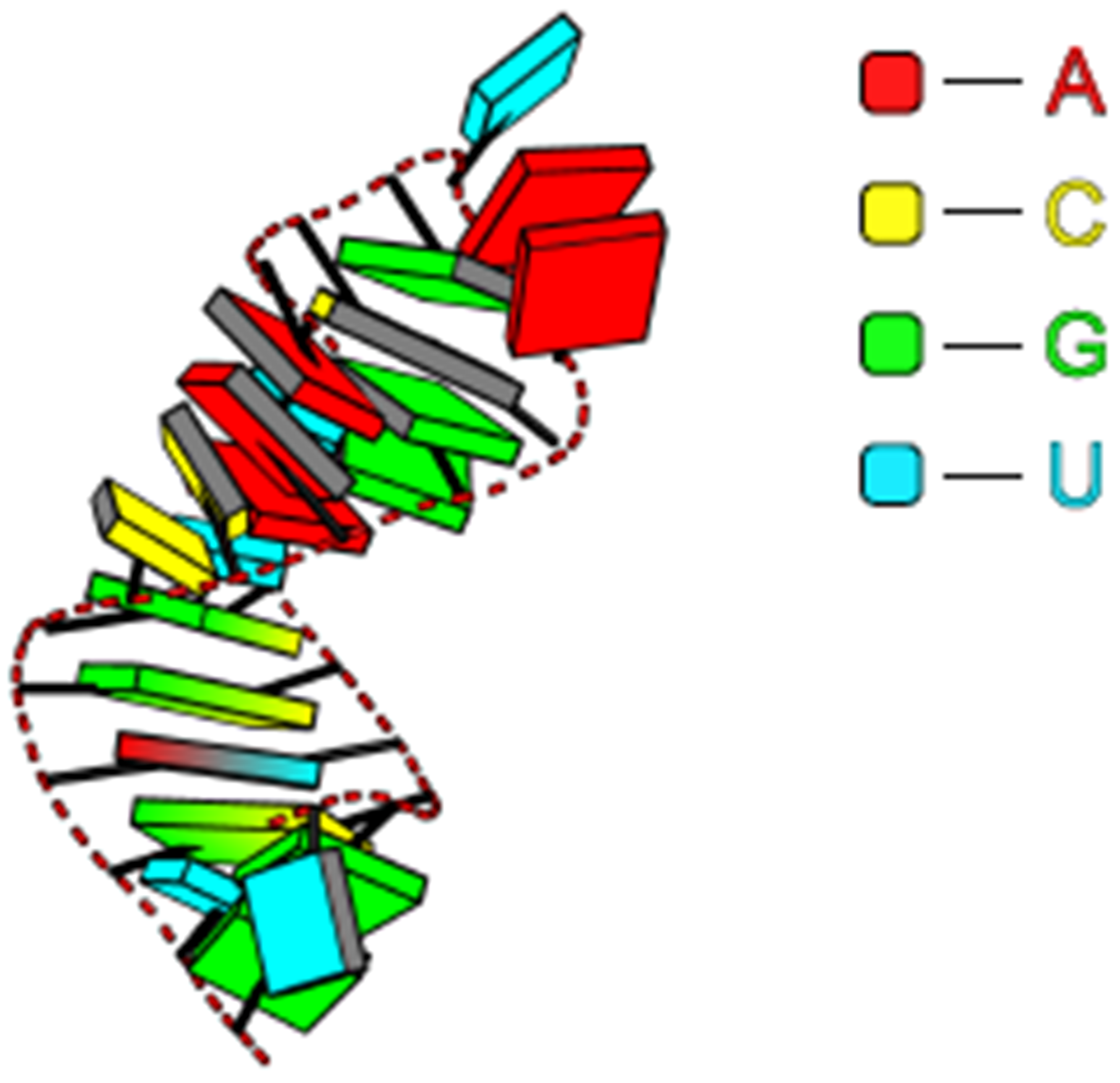
Nucleotide frames and canonical base pairs of 1MSY in the molecular dynamics trajectory. Canonical base pairs are drawn as single blocks. Minor edge grooves are displayed as gray edges of the rigid frames.

The analysis was performed using the following command:

~~~
dssna -f 1msy.md_npt.xtc -s 1msy.md_npt.tpr -sel RNA -o 1msy.nf.dat -hbo 1msy.hb.dat
-bpfo 1msy.bpf.dat -bpoo 1msy.bpo.dat -bpmo 1msy.bpm.dat -bso 1msy.bs.dat -bsoo 1msy.bso.dat
-bsmo 1msy.bsm.dat -ho 1msy.h.dat -dbno 1msy.dbn.dat -bpso 1msy.bps.dat -mmo 1msy.mmo.dat
-bbo 1msy.bb.dat -nospn -nohidebss -nochain
~~~

The command was executed on a server equipped with an Intel Xeon CPU E5-2660 v2 processor operating at 2.20 GHz. The trajectory analysis took 4 h 41 min 12.7 s. The total size of the output files was 40.2 GiB.

## Discussion

Analysis of the MD trajectory of the GUAA tetraloop mutant 1MSY revealed a substantial change in the overall RNA structure (see Figure 2). To analyze this transition in greater detail, we examined several key metrics. First, we analyzed the hydrogen-bonding networks formed between base pairs (see Figure S22). The number of hydrogen bonds began to decrease after approximately 200 ns, reaching a minimum at around 400 ns. Subsequently, the number of hydrogen bonds increased and eventually stabilized. The median number of hydrogen bonds over the entire MD trajectory was 16. We hypothesize that these observations indicate that the initial structure was in a metastable state and underwent a substantial conformational rearrangement to reach a new stable configuration.

**Figure 2.**
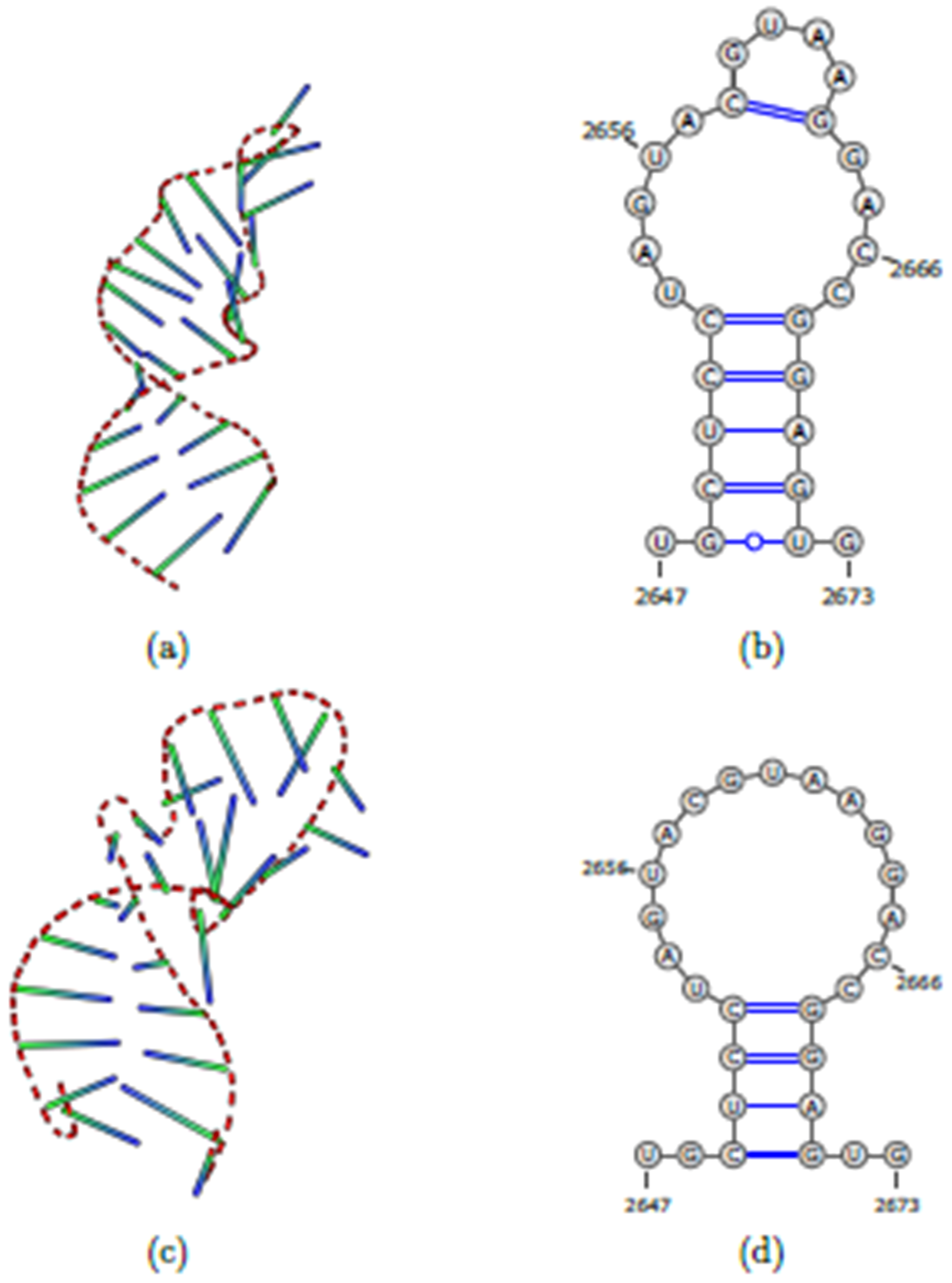
(2a) GUAA tetraloop mutant 1MSY at the beginning of the MD simulation. (2b) Secondary structure of the GUAA tetraloop mutant 1MSY at the beginning of the MD simulation, generated using VARNA[4]. (2c) GUAA tetraloop mutant 1MSY at the end of the MD simulation. (2d) Secondary structure of the GUAA tetraloop mutant 1MSY at the end of the MD simulation, generated using VARNA[4].

We also analyzed heat maps of base-pair occupancy and most stable lifetimes (see Figure S23). For completeness, we included all base-pair types, not only canonical ones. The maps revealed that the most highly populated base pairs were G2648*−* U2672, C2649 *−*G2671, U2650*−* A2670, C2651*−* G2669, C2652*−* G2668, and C2658*−* G2663. As expected, all of the most populated base pairs were canonical. Among these, the C2658 *−*G2663 pair exhibited the longest lifetime, stably forming an isolated base pair, but only at the beginning of the MD trajectory. The most persistent conformation accounted for slightly more than 3% of the total MD trajectory duration, indicating that base-pair formation was generally unstable. All non-canonical base pairs were found to be unstable, and the wobble pair G2648*−* U2672 was less stable than the Watson–Crick pairs.

We also analyzed base-stacking interactions in the GUAA tetraloop mutant 1MSY (see Figure S24). Notably, the time dependence of the number of base-stacking interactions closely resembles that of the hydrogen-bond count. The structure initially exhibited a high number of base-stacking interactions, which began to decrease after 200 ns, reached a minimum at approximately 400 ns, and subsequently stabilized. The median number of base-stacking interactions was 11. Since base-stacking interactions are not directly coupled to hydrogen bonds but are coordinated with them, we propose that their behavior supports our hypothesis that the GUAA tetraloop mutant 1MSY underwent a conformational transition from an initial metastable state.

We also analyzed heat maps of occupancy and most stable lifetimes for base-stacking interactions (see Figure S25). The occupancy map shows that stacking interactions most frequently form between adjacent nucleotides, as expected. The 3’-end exhibits higher base-stacking occupancy than the 5’-end. The most long-lived base-stacking interactions were observed between nucleotides G2648—C2649, A2657—A2665, G2663— G2664, G2668—G2669, and G2669—A2670. However, even the most stable stacking conformations persisted for no more than 0.20% of the total trajectory duration, which is a relatively small fraction. This indicates that base-stacking interactions are generally unstable, consistent with the overall structural instability of the system.

The backbone of the studied single-stranded RNA was highly flexible. The most mobile regions, in terms of bend formation, were G2655—U2656 (bends observed in 99.19% of the trajectory), U2660— A2661 (88.17%), G2664—A2665 (46.70%), A2662—G2663 (46.44%), G2648—C2649 (43.96%), U2653— A2654 (26.85%), U2672—G2673 (24%), A2657—C2658 (14.60%), and U2656—A2657 (13.36%). As expected, no chain breaks were detected.

The structure typically contained between 3 and 6 canonical base pairs. On average, one helix and one stem were present, while isolated base pairs were not observed (the C2658 *−*G2663 pair, which would be the most likely candidate for an isolated base pair, was too unstable after 200 ns). Four base pairs formed a stable stem throughout the trajectory: C2649*−* G2671, U2650*−* A2670, C2651 *−*G2669, and C2652 *−*G2668. This stable segment of canonical base pairs serves as the core around which the stable helix is formed (see Figure S26). At the beginning of the trajectory, 1MSY consistently formed two helical vectors, but after the destabilization of base pairs, it stably adopted only one, consistent with the observations described above. The secondary structure also underwent significant changes: the original structure contained one internal loop and one bulge (see Figure 2b), but after destabilization of the C2658 *−*G2663 isolated pair, the remaining internal loop was transformed into a hairpin (see Figure 2d).

The GUAA tetraloop mutant 1MSY is known to contain an A-minor motif (type X) formed by A2665 | G2655+U2656[2]. During the MD simulation, no A-minor motifs involving other nucleotides were observed. The aforementioned A-minor motif retained its type X classification throughout the trajectory. However, this A-minor motif exhibited high instability: the frequency of its formation decreased sharply after 100 ns. The most persistent occurrence of this motif lasted only 24 ps. The motif was most frequently formed at the beginning of the MD trajectory and was absent for 98% of the total simulation time. This instability likely results from the unstable formation of the G2655+U2656 base pair, as noted above.

## Conclusions

The DSSNA v2026 algorithm was applied to the analysis of a molecular dynamics trajectory of a GUAA tetraloop mutant of the sarcin–ricin domain of *Escherichia coli* 23S ribosomal RNA. The analysis showed that neither the base-pairing network nor the set of stacking interactions remains stable throughout the entire trajectory. Instead, only a subset of these interactions persists over extended periods. The most stable base pairs were identified, and all of them were canonical. Their persistence provides a structural framework that maintains the secondary structure of 1MSY and therefore constitutes its structural core.

The base-pairing profile indicates that the 1MSY structure undergoes substantial changes in both hydrogen-bonding and base-stacking interaction patterns during the MD trajectory. The structure provides a favorable environment for the formation of only one helix, which forms a stem through canonical base pairs. The isolated base pair C2658 *−*G2663 and the A-minor motif A2665 | G2655+U2656 were stable only at the beginning of the trajectory and disappeared after the hydrogen bonds and base-stacking interactions stabilized. The high flexibility of the sugar-phosphate backbone may impose geometric constraints on the formation and maintenance of stable base pairs. The secondary structure is determined primarily by the stem, not the isolated base pair. The disappearance of the C2658 *−*G2663 isolated base pair causes a change in the secondary structure profile of 1MSY.

The analysis of the 1MSY trajectory demonstrates that DSSNA can be used not only to identify individual structural elements but also to characterize their persistence and temporal behavior. By distinguishing stable interactions from transient ones, DSSNA provides a more detailed description of nucleic acid conformational dynamics than can be obtained from the analysis of a single static structure. This functionality may be particularly useful for studying the formation, disruption, and rearrangement of secondary and tertiary structural motifs in molecular dynamics simulations.

The present study is based on a single molecular dynamics trajectory and is intended primarily as a proof-of-concept application of DSSNA. Further validation using nucleic acids with different sequences, topologies, and conformational states will be required to assess the general applicability of the algorithm. Future development will focus on expanding the range of detectable structural motifs and improving the analysis of interactions in nucleic acids.

## Abbreviations

DSSNA: Define Spatial Structure of Nucleic Acids
MD: molecular dynamics
PDB: Protein Data Bank

## Data availability

The standalone version of DSSNA v2026 is freely available for download, installation, and use, without registration or a copy request, at https://gitlab.com/bio-pnpi/gmx-dssna. High-quality figures illustrating the algorithm, along with trajectory fragments and the corresponding analysis results, are publicly available at https://gitlab.com/bio-pnpi/gmx-dssna-refdata. Suggestions, questions, and bug reports are welcome via the project issue tracker or by contacting the authors.

## Supporting Information

Detailed glossary of terms describing the secondary and tertiary structure of nucleic acids, detailed description of the developed algorithm implementation in terms of the software algorithms, molecular dynamics simulation protocol for the GUAA tetraloop mutant of the sarcin–ricin ribosomal RNA domain, and additional graphs and figures visualizing the analysis results (PDF).

## Acknowledgments

Molecular visualizations were generated using the PyMOL Molecular Graphics System, Version 3.0.0 (Schrödinger, LLC)[5]. Schematic figures were prepared using Inkscape, version 1.4.3[6]. We thank the developers of these open-source and freely available tools for enabling the visualization of structural data.

The results of the work were obtained using computational resources of the supercomputing center of Peter the Great Saint-Petersburg Polytechnic University (https://www.spbstu.ru).

This work was supported by the Ministry of Science and Higher Education of the Russian Federation, grant No. 075-15-2024-630 (algorithm development) and topic 1024011000015-3-1.6.7 “Development of new computational methods in interdisciplinary studies of the structural and functional properties of biomacromolecules and biomacromolecular systems” (molecular dynamics and algorithm testing).

The authors declare no competing financial interest.

