## supporting information for "DSSNA: Automated Nucleic Acid Structure Analysis in Molecular Dynamics"

S. V. Gorelov[1,2,3], A. L. Konevega[1,2,3], and A. V. Shvetsov[1,2,3]

[1]Petersburg Nuclear Physics Institute named by B.P. Konstantinov of National Research Centre “Kurchatov Institute”, Gatchina 188300, Russia
[2]Peter the Great St. Petersburg Polytechnic University, St. Petersburg 195251, Russia
[3]National Research Centre “Kurchatov Institute”, Moscow 123098, Russia

* (S. V. G.), (A. L. K.), (A. V. S.)

#### Glossary

S.I A nucleotide frame (or nucleoframe) is a data structure that combines information about a nucleotide and its constituent atoms with information about the spatial arrangement of its base within a structure or molecular dynamics trajectory at a particular point in time (see Figure S1). Strictly speaking, a nucleoframe represents a nucleotide-specific coordinate reference frame[1]. Since the spatial arrangement of nucleotides may change over time, nucleoframes are recalculated at the beginning of each new trajectory frame. Thus, each new trajectory frame corresponds to a new nucleoframe for the same nucleotide.

Each nucleoframe is associated with a nucleotide-specific, right-handed Cartesian coordinate system. The orthonormal basis of this coordinate system is referred to as the frame (see S.II), whereas its origin is referred to as the origin (see S.III). Together, the frame and the origin define a unique nucleotide-specific reference frame. In addition to describing the relative spatial arrangement of nucleotides, nucleoframes make it possible to distinguish between different nucleotide edges, including the Watson–Crick edge, the Hoogsteen edge, the minor-groove edge, and the major-groove edge.

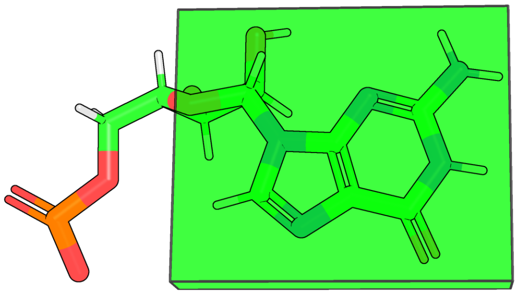

Figure S1: Visualization of a guanine nucleotide frame.

S.II The frame of a nucleotide is a right-handed orthonormal basis in affine space, consisting of a triad $\mathbf{T}$ (S1) of linearly independent orthonormal vectors $\widehat{\mathbf{x}}$, $\widehat{\mathbf{y}}$, and $\widehat{\mathbf{z}}$ attached to the nucleotide base.

$$\mathbf{T} = \begin{pmatrix} x_x & y_x & z_x \\ x_y & y_y & z_y \\ x_z & y_z & z_z \end{pmatrix} \tag{S1}$$

The frame is required to describe the relative three-dimensional spatial arrangement of nucleobases and base pairs. The vector $\widehat{\mathbf{x}}$ points toward the major groove. The vector $\widehat{\mathbf{y}}$ points toward the sugar-phosphate backbone, specifically toward the next nucleotide in the sequence. It is also parallel to the line connecting the C1' atom of the nucleotide of interest with the C1' atom of its idealized complementary nucleotide. For right-handed A-DNA and B-DNA conformations, the vector $\widehat{\mathbf{z}}$ is oriented in the 5'-to-3' direction of the sequence. The normal to the nucleotide frame, represented by the vector $\widehat{\mathbf{z}}$, is the same as the normal to the nucleotide base.

S.III The origin (O) of a nucleotide frame is a point in three-dimensional space that serves as the coordinate origin for the reference frame associated with a particular nucleotide base. In the standard reference frames[2], the positions of the origins are initially calculated by taking into account the inclination and separation of the complementary bases. The origin is defined as the midpoint between the Watson–Crick edges of an idealized complementary base pair.

In the standardized model, the origin is located at the intersection of the $\widehat{\mathbf{x}}$ axis of the base frame and the line connecting the C6 or C8 atom of one nucleotide to the corresponding C6 or C8 atom of the other nucleotide. The C6 atom is used for pyrimidines, whereas the C8 atom is used for purines.

S.IV The nucleoframe is calculated using the least-squares method. For each nucleotide in the structure, a minimal set of base atoms is selected sequentially and used to determine the base type[2]. A standardized base structure of the corresponding type (see Figure S2) is then selected in its minimum-energy conformation.

The coordinates of the corresponding atoms in the input and standardized structures are used to calculate their respective centroids. A covariance matrix is subsequently constructed from the centered atomic coordinates. The Davenport matrix is then derived from the covariance matrix, and the eigenvector corresponding to the largest eigenvalue is selected. This eigenvector is used to construct a Rodrigues' rotation matrix, which defines the basis of the nucleoframe.

The origin of the nucleoframe is calculated as the difference between the centroid of the base in the input structure and the centroid of the standardized base after the nucleoframe basis has been applied.

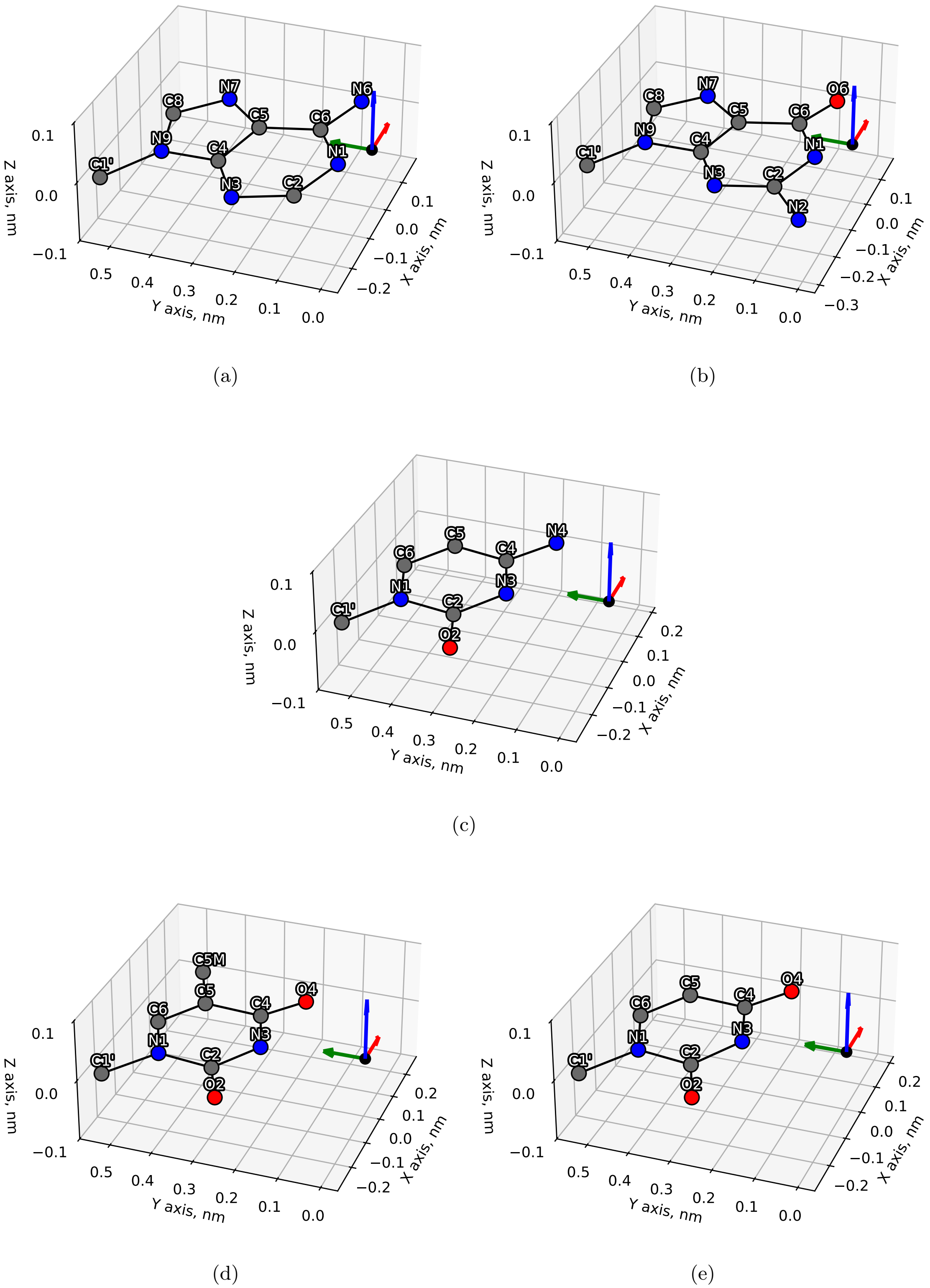


Figure S2: Standard reference frames[2] for adenine (S2a), guanine (S2b), cytosine (S2c), thymine (S2d) and uracil (S2e). The origin of the reference system is represented by a black dot.

S.V A hydrogen bond (HB) is an interaction between an electronegative atom, referred to as the hydrogen-bond acceptor, and a hydrogen atom that is covalently bonded to another electronegative atom, referred to as the hydrogen-bond donor (see Figure S3). Hydrogen-bond networks form secondary-structure patterns of varying complexity. They also play an important role in stabilizing nucleic acid helices.

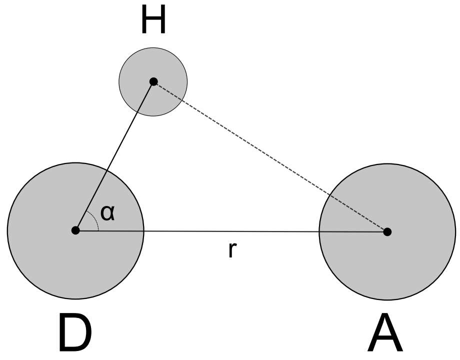


Figure S3: A hydrogen bond diagram. It involves a hydrogen-bond donor (D), a hydrogen-bond acceptor (A), and a hydrogen atom (H).

The hydrogen-bond detection algorithm implemented in DSSNA consists of two stages. The first stage involves the identification of hydrogen-bond donors and acceptors (see S.VI) and is performed before the first frame of a structure or molecular dynamics trajectory is analyzed. The second stage involves the direct detection of hydrogen bonds and is performed for each frame of a molecular dynamics trajectory (see S.VII).

S.VI The algorithm for identifying hydrogen-bond donors and acceptors is almost identical to that used in the `gmx hbond` algorithm[3], with only two exceptions:

1 A charge cutoff for acceptors is applied. The partial charge of an atom (in units of the elementary charge) must be more negative than a specified cutoff (the default value is $-0.05\ e$).

2 Only oxygen and nitrogen atoms can be acceptors. Hydrogen-bond donors are always nitrogen or oxygen atoms covalently bonded to hydrogen atoms.

S.VII At the start of each new frame of the molecular dynamics trajectory, a hydrogen bond search is performed. The algorithm uses a nearest-neighbor search method to identify potential hydrogen bonds between previously identified donors and acceptors.

To determine hydrogen bonds in trajectories, a geometric criterion for the existence of a hydrogen bond is used (S2), where $r$ is the distance between the hydrogen bond donor and acceptor, and $\alpha$ is the hydrogen-donor-acceptor angle (see Figure S3). The value $r_{HB} = 0.35$ nm corresponds to the first minimum of the radial distribution functions of water in the SPC/E model[4]. A hydrogen bond is considered to exist between the donor-acceptor pair if neither the angle nor the hydrogen-bond length exceeds the set cutoff. Both distance and angle cutoffs can be changed by the user if desired.

$$\begin{aligned} r &\leq r_{HB} = 0.35 \text{ nm} \\ \alpha &\leq \alpha_{HB} = 30^\circ \end{aligned} \tag{S2}$$

Although, according to the geometric criterion above, a hydrogen atom can theoretically form multiple hydrogen bonds in a system, DSSNA considers a hydrogen atom to be capable of participating in only one hydrogen bond. Therefore, only those hydrogen bonds with the lowest donor-acceptor distances $r$ and angles $\alpha$ (with a preference for angles) will be selected in the final set.

S.VIII A base pair is a fundamental structural unit of nucleic acids, consisting of two paired nucleotides that are connected by hydrogen bonds. Like nucleotides, base pairs also have individual frames (see Figure

S4). Within the DSSNA framework, a base pair is defined as a relationship between two nucleotides and exists only for a single trajectory frame.

For two nucleotides to be classified as a base pair, they must form at least one hydrogen bond between atoms belonging to the two nucleotides. Pairs for which no hydrogen bond is detected between the corresponding nucleobases are classified as false pairs or pseudo-pairs. By default, such pairs are excluded from the output statistics, although they may be used by internal algorithms.

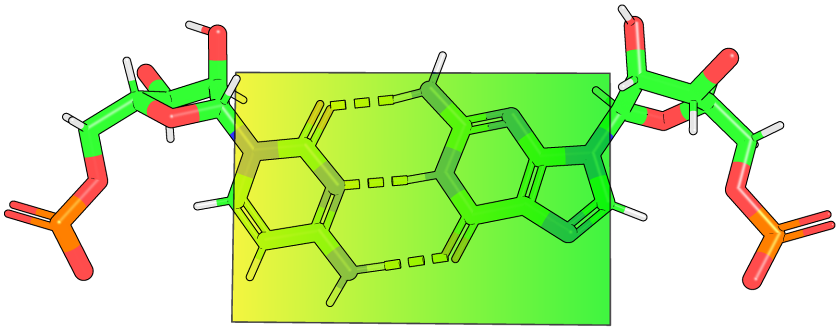

Figure S4: Visualization of a cytosine-guanine base pair (with frame).

In DSSNA, base pairs are denoted by two nucleotides, $N_1$ and $N_2$, separated by the "+", "-", or "±" symbols. The "+" symbol is used when the scalar product of the normals of the nucleotides in a base pair has a positive sign (they point in the same direction), the "-" symbol is used when the scalar product of the normals has a negative sign (they point in opposite directions), and the "±" symbol is used if the relative position of the normals cannot be determined for some reason or is meaningless. The notation "N±M" is equivalent to the notation "M±N". In the notation of base pairs, the nucleotide on the left is always the one that appears first in the notation of the structure (closer to the 5' end).

S.IX Two different nucleotides are considered to form a base pair if the following criteria are met:

1. The two nucleotides share at least one hydrogen bond;
2. The distance between the nucleotide origins does not exceed 1.5 nm;
3. The vertical separation (see S.X) between the nucleotides does not exceed 0.25 nm;
4. The angle between the nucleotide normals, taken modulo 180°, does not exceed 65°;
5. There are no base-stacking interactions (see S.XII) between the nucleotide bases of the potential pair.

S.X Vertical separation $d_s$ is a geometric characteristic of the projection of a point in space onto the geometry of a nucleotide. To determine vertical separation, we use the nucleotide origin $\vec{\mathbf{O}}_1$, its normal $\widehat{\mathbf{n}}$, and an arbitrary point in space $\vec{\mathbf{O}}_2$ for which we need to determine the vertical separation (usually this is either the origin of another nucleotide or the position of a capping atom). Mathematically, this is the scalar product of the normal and the vector from $\vec{\mathbf{O}}_1$ to $\vec{\mathbf{O}}_2$ (S3).

$$d_s = \widehat{\mathbf{n}} \cdot (\vec{\mathbf{O}}_2 - \vec{\mathbf{O}}_1). \tag{S3}$$

S.XI Calculating the angle $\alpha$ between vectors $\vec{\mathbf{a}}$ and $\vec{\mathbf{b}}$, taking into account the projection (most often between normals to nucleoframes or base pairs), is mathematically similar to calculating the angle between vectors, except that the resulting angle sign takes into account the relative direction of the vectors (S4). Thus, such an angle between codirectional vectors will be positive, and between oppositely directed vectors, negative.

$$\alpha = \cos^{-1}\left(\frac{\vec{\mathbf{a}} \cdot \vec{\mathbf{b}}}{\|\vec{\mathbf{a}}\| \cdot \|\vec{\mathbf{b}}\|}\right) \cdot \operatorname{sgn}(\vec{\mathbf{a}} \cdot \vec{\mathbf{b}}) = \cos^{-1}(\widehat{\mathbf{a}} \cdot \widehat{\mathbf{b}}) \cdot \operatorname{sgn}(\vec{\mathbf{a}} \cdot \vec{\mathbf{b}}). \tag{S4}$$

S.XII Base-stacking interactions (see Figure S5) are noncovalent attractive interactions between the aromatic surfaces of nucleobases arranged one above another in a stack. Together with hydrogen bonds between complementary base pairs, they contribute to the stabilization of nucleic acid helices. Base-stacking interactions arise from a combination of dispersion, electrostatic, and hydrophobic effects and promote the close and stable packing of nucleobases, thereby increasing the overall structural stability of nucleic acids. These interactions may occur between nucleobases in both DNA and RNA and can substantially influence the stabilization of their three-dimensional organization.

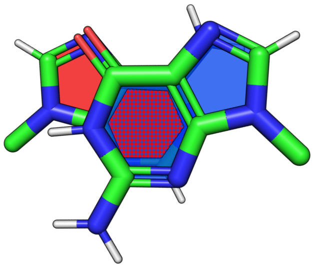

Figure S5: Visualization of the stacking area between guanine and adenine bases.

An ordered set of nucleotides assembled through base-stacking interactions, irrespective of whether the corresponding nucleotides are connected within the sugar-phosphate backbone, defines a base stack. In DSSNA, base-stacking interactions between two nucleobases are identified using either an assessment of base overlap (see S.XIII) or an analysis of the geometric parameters of the corresponding nucleotides (see S.XIV).

S.XIII The base overlap estimation method is the default method used in DSSNA. It involves directly estimating the area of overlap between the aromatic rings of two nucleobases (see Figure S6).

Its criteria are as follows:

1. The angle $\alpha$ between the two nucleobase normals, taking into account the projection sign (see S.XI), does not exceed 23° (the value can be varied by the user);
2. The ratio of the overlap area of the bases of two nucleotides projected onto the same plane to the area of the smaller of the two bases exceeds the set threshold of 10% (the value can be varied by the user).

In more detail, the algorithm for estimating the overlap area of the bases of two nucleobases is as follows:

1. The average origin $\vec{\mathbf{O}}_m$ of two nucleobases is found (S5).

$$\vec{\mathbf{O}}_m = \frac{\vec{\mathbf{O}}_1 + \vec{\mathbf{O}}_2}{2} \tag{S5}$$

2. The average vector of two normals of the compared nucleobases is calculated (S6).

$$\widehat{\mathbf{n}}_m = \frac{\widehat{\mathbf{n}}_1 + \widehat{\mathbf{n}}_2}{2} \tag{S6}$$

3. The aromatic ring atoms at nucleobases 1 and 2 are projected onto a common plane and transferred (rotationally) to the XY plane (from the average normal to the vector $\widehat{\mathbf{k}} = \{0, 0, 1\}$), so that the Z component does not play a role in the calculation;
4. Two polygons are formed, the vertices of which are the aromatic ring atoms of nucleobases 1 and 2 projected onto the XY plane;
5. Using polygon clipping, the Sutherland–Hodgman algorithm trims the surface where two polygons overlap (if any);
6. The area of the resulting clipped polygon is calculated using the Gaussian formula.
7. The ratio of the resulting base overlap area to the smaller of the two nucleotide base areas is calculated. The value, by definition, is always in the range $[0, 1]$ and is subsequently used for comparison with the cutoff value (0.1 or 10% by default), which can be varied by the user. Normalization to the area of the smaller base has a clear interpretation: it shows what proportion of the smaller base is involved in the overlap. To assess the sensitivity of base-stacking detection to the overlap threshold, analyses were performed using overlap cutoffs of 5%, 10%, and 20%. The 10% threshold was selected as the default value because it provided a balance between detecting partially overlapping stacked bases and limiting spurious contacts.

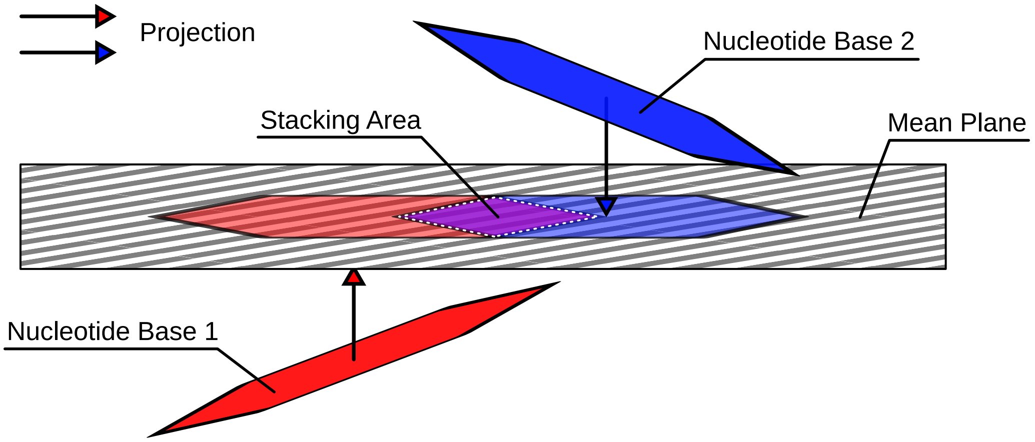


Figure S6: Diagram describing the method for assessing the overlap of nucleobases.

S.XIV The method for assessing base-stacking interactions based on the analysis of geometric parameters of nucleobases (see Figure S7) is a relatively simple method that consists of calculating several geometric parameters between two nucleobases and checking whether they are within an acceptable range.

The criteria for assessing the existence of a base-stacking conformation in this method are as follows:

1. The angle $\alpha$ between two nucleobase normals, taking into account the projection sign (see S.XI), does not exceed 23° (the value can be varied by the user);
2. There is at least one vector $\vec{\mathbf{d}}_{12}$ (or $\vec{\mathbf{d}}_{21}$) passing from the geometric center of any aromatic ring at nucleobase 1 to the geometric center of any aromatic ring at nucleobase 2, the length of which does not exceed 0.45 nm (the value can be varied by the user);
3. The angle $\beta$ between $\vec{\mathbf{d}}_{12}$ and $\widehat{\mathbf{n}}_1$ (or between $\vec{\mathbf{d}}_{21}$ and $\widehat{\mathbf{n}}_2$) does not exceed 40° (the value can be varied by the user).

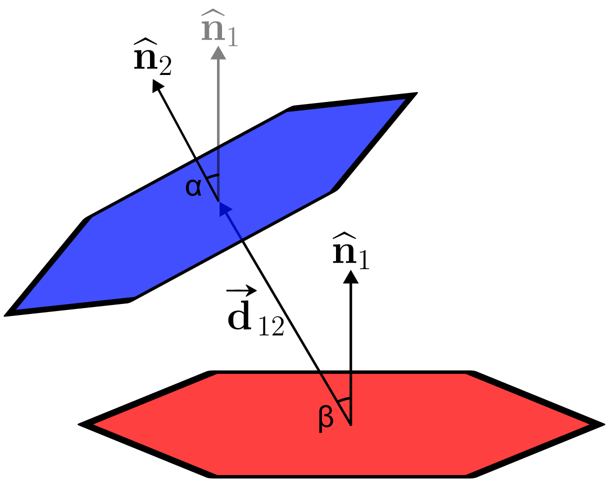


Figure S7: Diagram describing the method for assessing base-stacking interactions based on the analysis of geometric parameters of nucleobases.

S.XV In DSSNA, base pairs can have up to 9 different types (see Table S1). The geometric parameters of the formed base pair, the types of paired bases, and, in some cases, hydrogen bond networks play a decisive role in determining base pair types in DSSNA. Of all the base pair types defined in DSSNA, only Watson–Crick ($WC$) and Wobble ($W$) pairs are canonical. Canonical base pairs play a key role in determining stems, isolated pairs and kissing loops.

Table S1: Base pair types and their designations in DSSNA.

| **Base Pair Type** | **Designation** | **Criterion** |
|---|---|---|
| Unspecified | – | |
| Watson–Crick | $WC$ | Complementary R−Y with acceptable geometry |
| Reverse Watson–Crick | $RWC$ | Complementary R+Y with acceptable geometry |
| Hoogsteen | $H$ | R+Y with acceptable geometry |
| Reverse Hoogsteen | $RH$ | R−Y with acceptable geometry |
| Wobble | $W$ | G−U with acceptable geometry |
| Sheared[5] | $S$ | G−A with N2-H—N7 and N6-H—N3 H-bonds |
| Calcutta[6, 7] | $C$ | U−U with N3-H—O4 H-bond and at least one C-H—O H-bond |
| Platform[8] | $P$ | Adjacent N±N with at least one base-base H-bond |

Acceptable geometry means that the deviation of the corresponding base vectors of two nucleotides is within the acceptable range. In DSSNA, acceptable Watson–Crick, Wobble, and Hoogsteen base pair geometry for two nucleotides $N_1$ and $N_2$ implies mandatory directionality of the vectors $\widehat{\mathbf{z}}_1$ and $\widehat{\mathbf{z}}_2$ to a certain degree of accuracy specified by the cutoff $\theta$ (the value can be varied by the user). For R+Y pairs, the vectors $\widehat{\mathbf{z}}_1$ and $\widehat{\mathbf{z}}_2$ must be parallel (S7).

$$\cos^{-1}(\widehat{\mathbf{z}}_1 \cdot \widehat{\mathbf{z}}_2)\frac{180}{\pi} \in [0^\circ; \theta] \tag{S7}$$

whereas for pairs of type R−Y they must be antiparallel (S8).

$$\cos^{-1}(\widehat{\mathbf{z}}_1 \cdot \widehat{\mathbf{z}}_2)\frac{180}{\pi} \in [180^\circ - \theta; 180^\circ] \tag{S8}$$

For Watson–Crick and Wobble pairs, it is additionally required that the vectors $\widehat{\mathbf{x}}_1$ and $\widehat{\mathbf{x}}_2$ be parallel, and the vectors $\widehat{\mathbf{y}}_1$ and $\widehat{\mathbf{y}}_2$ be antiparallel (S9).

$$\begin{cases} \cos^{-1}(\widehat{\mathbf{x}}_1 \cdot \widehat{\mathbf{x}}_2)\frac{180}{\pi} \in [0°; \theta] \\ \cos^{-1}(\widehat{\mathbf{y}}_1 \cdot \widehat{\mathbf{y}}_2)\frac{180}{\pi} \in [180° - \theta; 180°] \end{cases} \tag{S9}$$

For Hoogsteen pairs, an additional requirement is that there is an angle of 90° between the vectors $\widehat{\mathbf{x}}_1$ and $\widehat{\mathbf{x}}_2$ and between the vectors $\widehat{\mathbf{y}}_1$ and $\widehat{\mathbf{y}}_2$ (S10).

$$\begin{cases} \cos^{-1}(\widehat{\mathbf{x}}_1 \cdot \widehat{\mathbf{x}}_2)\frac{180}{\pi} \in [90° - \frac{\theta}{2}; 90° + \frac{\theta}{2}] \\ \cos^{-1}(\widehat{\mathbf{y}}_1 \cdot \widehat{\mathbf{y}}_2)\frac{180}{\pi} \in [90° - \frac{\theta}{2}; 90° + \frac{\theta}{2}] \end{cases} \tag{S10}$$

S.XVI Base pairs have three types of orientation relative to each other (see Table S2). The orientation type depends on the scalar product of the normal of one base $\widehat{\mathbf{n}}_1$ and the normal of the second base $\widehat{\mathbf{n}}_2$.

Table S2: Supported base pair orientations.

| **Base Pair Orientation** | **Designation** | **Criterion** |
|---|---|---|
| Unspecified | ± | |
| Default | − | $\widehat{\mathbf{n}}_1 \cdot \widehat{\mathbf{n}}_2 < 0$ |
| Flipped | + | $\widehat{\mathbf{n}}_1 \cdot \widehat{\mathbf{n}}_2 > 0$ |

S.XVII A virtual bond (A–B) is a vector connecting atom A of one nucleotide to atom B of another nucleotide, denoted by $\overrightarrow{\mathbf{AB}}$, where the two atoms are not connected by a covalent bond[9]. Virtual bonds are used as structural descriptors for characterizing the geometry of nucleic acids. In DSSNA, virtual bonds are defined both between consecutive nucleotides in the same strand and between the nucleotides forming a base pair.

In some output files, virtual-bond parameters are represented using notations such as (RA/YB). This notation indicates that the atoms defining the virtual bond depend on the nucleotide base type: R denotes a purine and Y denotes a pyrimidine. For purines, the virtual bond is defined using atom A, whereas for pyrimidines it is defined using atom B.

The output may also include angles between virtual bonds. In the primary output, $\lambda_1$ denotes the angle between the virtual bonds C1'–(RN9/YN1) and C1'–C1'. The angle $\lambda_2$ is defined between the virtual bonds (RN9/YN1)–C1' and C1'–C1'.

The algorithm determines the following parameters of virtual bonds:

1. Length of virtual bond C1'–C1';
2. Length of virtual bond (RN9/YN1)–(RN9/YN1);
3. Length of virtual bond (RC8/YC6)–(RC8/YC6);
4. Length of virtual bond P–P;
5. Angle $\lambda_1$ (for base pairs only);
6. Angle $\lambda_2$ (for base pairs only).

S.XVIII A dinucleotide step is a structural arrangement comprising two non-overlapping base pairs. Consider two nucleotides, $i$ and $j$, that are adjacent in the structure and are not separated by a break (see S.XLVII). If nucleotide $i$ forms a base pair (rather than a pseudo-pair) with nucleotide $k$, nucleotide $j$ forms a base pair (rather than a pseudo-pair) with nucleotide $l$, and nucleotides $k$ and $l$ are adjacent and not separated by a break, then nucleotides $i$, $j$, $k$, and $l$ are considered to form a dinucleotide step[10]. A helical step is analogous to a dinucleotide step, but is defined in a coordinate system aligned with the local helical axis rather than the base-pair frame.

Both dinucleotide and helical steps provide critical rigid-body parameters that describe the relative orientation of consecutive base pairs (see S.XXII, S.XXIII). These parameters can subsequently be employed to analyze and classify helical conformations (see S.XXIV), offering insights into the structural properties of nucleic acids.

S.XIX In some parts of the algorithm, it is necessary to rotate a triad of vectors by an angle $\theta$ about the axis $\widehat{\mathbf{k}} = \{k_x, k_y, k_z\}$. This is achieved by applying Rodrigues' rotation formula (S11).

$$\mathbf{R}_k(\theta) = \begin{bmatrix} c + (1-c)k_x^2 & (1-c)k_x k_y - s k_z & (1-c)k_x k_z + s k_y \\ (1-c)k_x k_y + s k_z & c + (1-c)k_y^2 & (1-c)k_y k_z - s k_x \\ (1-c)k_x k_z - s k_y & (1-c)k_y k_z + s k_x & c + (1-c)k_z^2 \end{bmatrix} \tag{S11}$$

where $s = \sin(\theta)$ and $c = \cos(\theta)$.

S.XX In the structural analysis of nucleic acids, considerable importance is attached to the characterization of the relative arrangement of the rigid frames associated with the nucleotide bases forming a base pair. Six standard rigid-body parameters are widely used for this purpose: three translational parameters describing relative displacement and three rotational parameters describing relative orientation (see Table S3).

Table S3: Six rigid-body parameters for nucleic acid structures.

| **Parameter** | **Base Pair** | **Dinucleotide Step** | **Helical Step** |
|---|---|---|---|
| translational (x-axis) | Shear | Shift | x-displacement |
| translational (y-axis) | Stretch | Slide | y-displacement |
| translational (z-axis) | Stagger | Rise | Helical Rise |
| rotational (x-axis) | Buckle | Tilt | Inclination |
| rotational (y-axis) | Propeller | Roll | Tip |
| rotational (z-axis) | Opening | Twist | Helical Twist |

S.XXI When a base pair is detected, the DSSNA algorithm constructs a base pair rigid frame. This is constructed from the rigid frames of the bases that form the pair. Like the rigid frame of a nucleotide, the base pair rigid frame has an origin and a basis. Like all frames, it has three translational and three rotational parameters, which the algorithm calculates upon building a frame (see Figure S8)[10].

The origin of the base pair is defined as the midpoint between the origins of the two bases (S12).

$$\vec{\mathbf{O}}_{bp} = \frac{\vec{\mathbf{O}}_1 + \vec{\mathbf{O}}_2}{2} \tag{S12}$$

Next, the algorithm attempts to determine the basis triad of base pair vectors. To do this, it uses the bases of the base pair's nucleotides. If the nucleotide normals are in opposite directions, the algorithm inverts the sign of the $\widehat{\mathbf{y}}_2$ and $\widehat{\mathbf{z}}_2$ vectors for the second nucleotide in the pair (closest to the 3' end).

Then the Buckle-Opening angle $\gamma$ is defined as the angle between the axes $\widehat{\mathbf{y}}_1$ and $\widehat{\mathbf{y}}_2$ (S13).

$$\gamma = \cos^{-1}(\widehat{\mathbf{y}}_1 \cdot \widehat{\mathbf{y}}_2)\frac{180}{\pi} \tag{S13}$$

The Buckle-Opening axis $\overrightarrow{\mathbf{bo}}$ is defined as the vector product of $\widehat{\mathbf{y}}_1$ and $\widehat{\mathbf{y}}_2$ (S14).

$$\overrightarrow{\mathbf{bo}} = \widehat{\mathbf{y}}_1 \times \widehat{\mathbf{y}}_2 \tag{S14}$$

$\overrightarrow{\mathbf{bo}}$ is normalized to $\widehat{\mathbf{bo}}$. Next, we rotate the basis of the first nucleotide base by an angle $-\frac{\gamma}{2}$ about the $\widehat{\mathbf{bo}}$ axis and the basis of the second nucleotide base by an angle $\frac{\gamma}{2}$ about the $\widehat{\mathbf{bo}}$ axis (S15).

$$\begin{aligned} \mathbf{T}_1' &= \mathbf{R}_{bo}(-\frac{\gamma}{2}) \cdot \mathbf{T}_1 \\ \mathbf{T}_2' &= \mathbf{R}_{bo}(\frac{\gamma}{2}) \cdot \mathbf{T}_2 \end{aligned} \tag{S15}$$

The basis triad of the base pair $\mathbf{T}_{bp}$, which contains the vectors $\widehat{\mathbf{x}}_{bp}$, $\widehat{\mathbf{y}}_{bp}$, and $\widehat{\mathbf{z}}_{bp}$, is obtained by averaging $\mathbf{T}'_1$ and $\mathbf{T}'_2$ (S16).

$$\begin{aligned}\widehat{\mathbf{x}}_{bp} &= \frac{\widehat{\mathbf{x}}'_1 + \widehat{\mathbf{x}}'_2}{2}\\ \widehat{\mathbf{y}}_{bp} &= \frac{\widehat{\mathbf{y}}'_1 + \widehat{\mathbf{y}}'_2}{2}\\ \widehat{\mathbf{z}}_{bp} &= \frac{\widehat{\mathbf{z}}'_1 + \widehat{\mathbf{z}}'_2}{2}\end{aligned} \tag{S16}$$

The vector $\widehat{\mathbf{z}}_{bp}$ is the base pair normal (S17).

$$\widehat{\mathbf{z}}_{bp} = \widehat{\mathbf{n}}_{bp} \tag{S17}$$

Propeller $\omega$ (see Figure S8e) — the angle between the transformed axes $\widehat{\mathbf{x}}'_1$ and $\widehat{\mathbf{x}}'_2$ (or $\widehat{\mathbf{z}}'_1$ and $\widehat{\mathbf{z}}'_2$) — is defined as follows (S18).

$$\omega = (\cos^{-1}(\widehat{\mathbf{x}}'_1 \cdot \widehat{\mathbf{x}}'_2)\frac{180}{\pi})(\mathrm{sgn}((\widehat{\mathbf{x}}'_2 \times \widehat{\mathbf{x}}'_1) \cdot \widehat{\mathbf{y}}_{bp})) \tag{S18}$$

The algorithm then determines the angle $\phi$, which is the angle between $\widehat{\mathbf{bo}}$ and $\widehat{\mathbf{x}}_{bp}$ (S19). Using this angle, we can calculate the Buckle $\kappa$ (S20) and Opening $\sigma$ (see Figure S8f).

$$\phi = (\cos^{-1}(\widehat{\mathbf{bo}} \cdot \widehat{\mathbf{x}}_{bp})\frac{180}{\pi})(\mathrm{sgn}((\widehat{\mathbf{x}}_{bp} \times \widehat{\mathbf{bo}}) \cdot \widehat{\mathbf{y}}_{bp})) \tag{S19}$$

$$\kappa = \gamma\cos(\phi) \tag{S20}$$

$$\sigma = \gamma\sin(\phi) \tag{S21}$$

Shear $S_x$ (see Figure S8a), Stretch $S_y$ (see Figure S8b), and Stagger $S_z$ (see Figure S8c) are defined as projections of the vector starting from $\vec{\mathbf{O}}_2$ and going to $\vec{\mathbf{O}}_1$ onto the corresponding vectors of the basis triad $\mathbf{T}_{bp}$ (S22).

$$\vec{\mathbf{S}} = (\vec{\mathbf{O}}_1 - \vec{\mathbf{O}}_2) \cdot \mathbf{T}_{bp} \tag{S22}$$

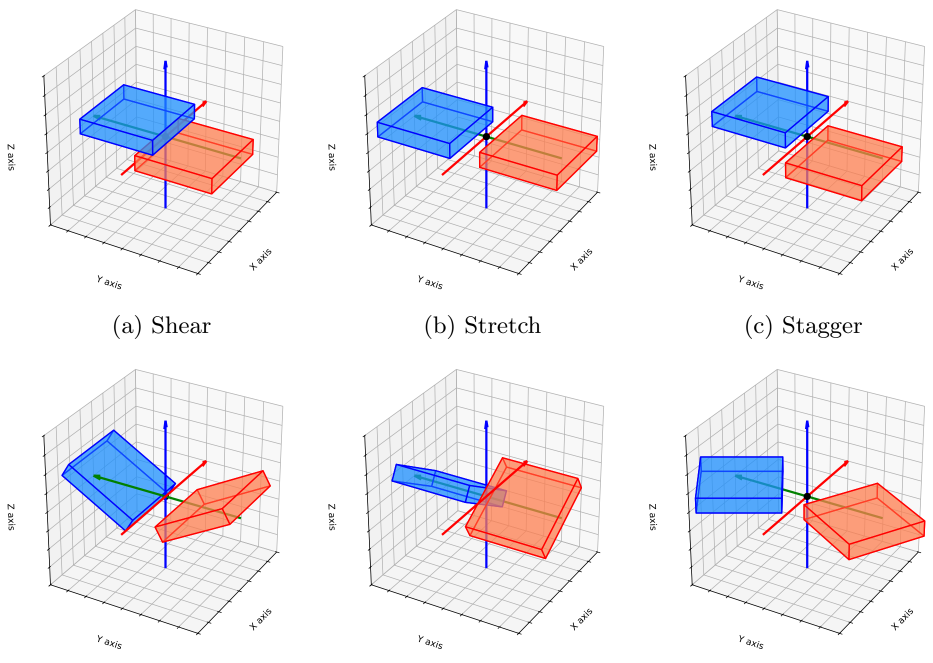


(a) Shear (b) Stretch (c) Stagger

(d) Buckle (e) Propeller (f) Opening

Figure S8: Six rigid-body parameters of base pairs.

S.XXII Upon finding a dinucleotide step, the DSSNA algorithm calculates the parameters of its rigid frame. This is constructed based on the rigid frames of the base pairs that comprise the dinucleotide step. The dinucleotide step also has three translational and three rotational parameters (see Figure S9)[10].

The origin of the dinucleotide step is defined as the midpoint between the base pair origins in the step (S23).

$$\vec{\mathbf{O}}_{ds} = \frac{\vec{\mathbf{O}}_{bp1} + \vec{\mathbf{O}}_{bp2}}{2} \tag{S23}$$

Next, an attempt is made to determine the basis triad of the dinucleotide step. This determination is made using the basis triads of the step's base pairs. If the base pair normals are in opposite directions, the algorithm inverts the sign of the vectors $\widehat{\mathbf{y}}_{bp2}$ and $\widehat{\mathbf{z}}_{bp2}$ for the second base pair in the step (closest to the 3' end).

Then the Roll-Tilt angle $\Gamma$ is defined as the angle between the axes $\widehat{\mathbf{z}}_{bp1}$ and $\widehat{\mathbf{z}}_{bp2}$ (S24).

$$\Gamma = \cos^{-1}(\widehat{\mathbf{z}}_{bp1} \cdot \widehat{\mathbf{z}}_{bp2}) \frac{180}{\pi} \tag{S24}$$

The Roll-Tilt axis $\vec{\mathbf{rt}}$ is defined as the vector product of $\widehat{\mathbf{z}}_{bp1}$ and $\widehat{\mathbf{z}}_{bp2}$ (S25).

$$\vec{\mathbf{rt}} = \widehat{\mathbf{z}}_{bp1} \times \widehat{\mathbf{z}}_{bp2} \tag{S25}$$

$\vec{\mathbf{rt}}$ is normalized, yielding $\widehat{\mathbf{rt}}$. Next, we rotate the basis of the first base pair by an angle $-\frac{\Gamma}{2}$ about the $\widehat{\mathbf{rt}}$ axis and the basis of the second base pair by an angle $\frac{\Gamma}{2}$ about the $\widehat{\mathbf{rt}}$ axis (S26).

$$\begin{aligned} \mathbf{T}'_{bp1} &= \mathbf{R}_{rt}(-\frac{\Gamma}{2}) \cdot \mathbf{T}_{bp1} \\ \mathbf{T}'_{bp2} &= \mathbf{R}_{rt}(\frac{\Gamma}{2}) \cdot \mathbf{T}_{bp2} \end{aligned} \tag{S26}$$

The basis triad of the dinucleotide step $\mathbf{T}_{ds}$, which contains the vectors $\widehat{\mathbf{x}}_{ds}$, $\widehat{\mathbf{y}}_{ds}$, and $\widehat{\mathbf{z}}_{ds}$, is obtained by averaging $\mathbf{T}'_{bp1}$ and $\mathbf{T}'_{bp2}$ (S27).

$$\begin{aligned}\widehat{\mathbf{x}}_{ds} &= \frac{\widehat{\mathbf{x}}'_{bp1} + \widehat{\mathbf{x}}'_{bp2}}{2} \\ \widehat{\mathbf{y}}_{ds} &= \frac{\widehat{\mathbf{y}}'_{bp1} + \widehat{\mathbf{y}}'_{bp2}}{2} \\ \widehat{\mathbf{z}}_{ds} &= \frac{\widehat{\mathbf{z}}'_{bp1} + \widehat{\mathbf{z}}'_{bp2}}{2}\end{aligned} \tag{S27}$$

Next, we determine Twist $\Omega$ (see Figure S9f) — the angle between the transformed axes $\widehat{\mathbf{y}}'_{bp1}$ and $\widehat{\mathbf{y}}'_{bp2}$ (or $\widehat{\mathbf{x}}'_{bp1}$ and $\widehat{\mathbf{x}}'_{bp2}$) taking into account the direction $\widehat{\mathbf{z}}_{ds}$ (S28).

$$\Omega = (\cos^{-1}(\widehat{\mathbf{y}}'_{bp1} \cdot \widehat{\mathbf{y}}'_{bp2})\frac{180}{\pi})(\mathrm{sgn}((\widehat{\mathbf{y}}'_{bp2} \times \widehat{\mathbf{y}}'_{bp1}) \cdot \widehat{\mathbf{z}}_{ds})) \tag{S28}$$

The algorithm determines the angle $\phi$, which is the angle between $\widehat{\mathbf{rt}}$ and $\widehat{\mathbf{y}}_{ds}$ (S29). Using this angle, we can calculate Roll $\rho$ (see Figure S9e) (S30) and Tilt $\tau$ (see Figure S9d) (S31).

$$\phi = (\cos^{-1}(\widehat{\mathbf{rt}} \cdot \widehat{\mathbf{y}}_{ds})\frac{180}{\pi})(\mathrm{sgn}((\widehat{\mathbf{y}}_{ds} \times \widehat{\mathbf{rt}}) \cdot \widehat{\mathbf{z}}_{ds})) \tag{S29}$$

$$\rho = \Gamma \cos(\phi) \tag{S30}$$

$$\tau = \Gamma \sin(\phi) \tag{S31}$$

Shift $D_x$ (see Figure S9a), Slide $D_y$ (see Figure S9b), and Rise $D_z$ (see Figure S9c) are defined as projections of the vector starting from $\vec{\mathbf{O}}_{bp1}$ and going to $\vec{\mathbf{O}}_{bp2}$ onto the corresponding vectors of the basis triad $\mathbf{T}_{ds}$ (S32).

$$\vec{\mathbf{D}} = (\vec{\mathbf{O}}_{bp2} - \vec{\mathbf{O}}_{bp1}) \cdot \mathbf{T}_{ds} \tag{S32}$$

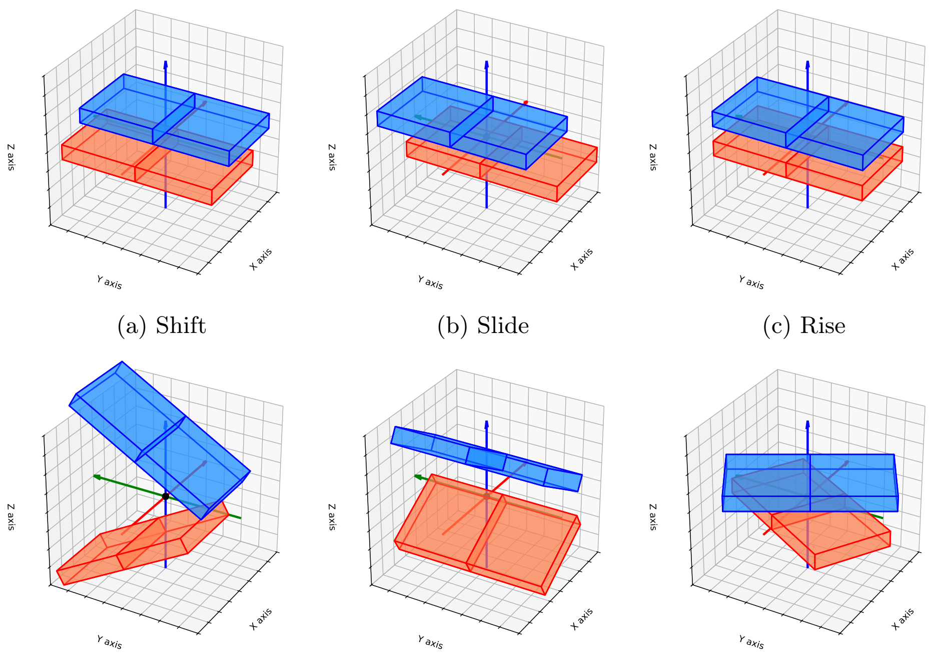


(a) Shift (b) Slide (c) Rise

(d) Tilt (e) Roll (f) Twist

Figure S9: Six rigid-body parameters of dinucleotide steps.

S.XXIII In addition to the rigid frame parameters of the dinucleotide step, the DSSNA algorithm can analyze the rigid frame parameters of the local helical step. These are constructed using the rigid frames of the base pairs that comprise the helical (and, accordingly, dinucleotide) step. The helical step also has three translational and three rotational parameters (see Figure S10)[10].

First, the basis triad of the helical step is determined. This determination is made using the basis triads of the base pairs in the step. If the base pair normals are in opposite directions, the algorithm inverts the sign of the vectors $\widehat{\mathbf{y}}_{bp2}$ and $\widehat{\mathbf{z}}_{bp2}$ for the second base pair in the step (closest to the 3' end).

Initially, the local helical axis $\overrightarrow{\mathbf{h}}$ is determined from the vectors $\widehat{\mathbf{x}}_{bp}$ and $\widehat{\mathbf{y}}_{bp}$ of the two base pairs in the step (S33).

$$\overrightarrow{\mathbf{h}} = (\widehat{\mathbf{x}}_{bp2} - \widehat{\mathbf{x}}_{bp1}) \times (\widehat{\mathbf{y}}_{bp2} - \widehat{\mathbf{y}}_{bp1}) \tag{S33}$$

The vector $\overrightarrow{\mathbf{h}}$ is normalized to give $\widehat{\mathbf{h}}$. The Tip-Inclination angle $\Psi$ is then defined as the angle between the axes $\widehat{\mathbf{h}}$ and $\widehat{\mathbf{z}}_{bp1}$ (S34).

$$\Psi = \cos^{-1}(\widehat{\mathbf{h}} \cdot \widehat{\mathbf{z}}_{bp1})\frac{180}{\pi} \tag{S34}$$

Next, the hinge axes are defined (S35).

$$\begin{aligned} \overrightarrow{\mathbf{H}}_1 &= \widehat{\mathbf{h}} \times \widehat{\mathbf{z}}_{bp1} \\ \overrightarrow{\mathbf{H}}_2 &= \widehat{\mathbf{h}} \times \widehat{\mathbf{z}}_{bp2} \end{aligned} \tag{S35}$$

The vectors $\overrightarrow{\mathbf{H}}_1$ and $\overrightarrow{\mathbf{H}}_2$ are normalized to give $\widehat{\mathbf{H}}_1$ and $\widehat{\mathbf{H}}_2$. The basis triads $\mathbf{T}_{bp1}$ and $\mathbf{T}_{bp2}$ rotate relative to the axes $\widehat{\mathbf{H}}_1$ and $\widehat{\mathbf{H}}_2$, respectively, by the angle $-\Psi$ (S36).

$$\begin{aligned}\mathbf{T}'_{hs1} &= \mathbf{R}_{H_1}(-\Psi)\cdot\mathbf{T}_{bp1}\\ \mathbf{T}'_{hs2} &= \mathbf{R}_{H_2}(-\Psi)\cdot\mathbf{T}_{bp2}\end{aligned} \tag{S36}$$

The basis triad of the helical step $\mathbf{T}_{hs}$, which contains the vectors $\widehat{\mathbf{x}}_{hs}$, $\widehat{\mathbf{y}}_{hs}$, and $\widehat{\mathbf{z}}_{hs}$, is obtained by averaging $\mathbf{T}'_{hs1}$ and $\mathbf{T}'_{hs2}$ (S37).

$$\begin{aligned}\widehat{\mathbf{x}}_{hs} &= \frac{\widehat{\mathbf{x}}'_{hs1}+\widehat{\mathbf{x}}'_{hs2}}{2}\\ \widehat{\mathbf{y}}_{hs} &= \frac{\widehat{\mathbf{y}}'_{hs1}+\widehat{\mathbf{y}}'_{hs2}}{2}\\ \widehat{\mathbf{z}}_{hs} &= \frac{\widehat{\mathbf{z}}'_{hs1}+\widehat{\mathbf{z}}'_{hs2}}{2}\end{aligned} \tag{S37}$$

Next, we define Helical Twist $\Omega$ (see Figure S10f) — the angle between the transformed axes $\widehat{\mathbf{y}}'_{hs1}$ and $\widehat{\mathbf{y}}'_{hs2}$ (or $\widehat{\mathbf{x}}'_{hs1}$ and $\widehat{\mathbf{x}}'_{hs2}$) taking into account the direction $\widehat{\mathbf{h}}$ (S38).

$$\Omega = (\cos^{-1}(\widehat{\mathbf{y}}'_{hs1}\cdot\widehat{\mathbf{y}}'_{hs2})\frac{180}{\pi})(\operatorname{sgn}((\widehat{\mathbf{y}}'_{hs2}\times\widehat{\mathbf{y}}'_{hs1})\cdot\widehat{\mathbf{h}})) \tag{S38}$$

Helical rise $d_z$ (see Figure S10c) is the projection of the vector starting from $\vec{\mathbf{O}}_{bp1}$ and going to $\vec{\mathbf{O}}_{bp2}$ onto the vector $\widehat{\mathbf{h}}$ (S39).

$$d_z = (\vec{\mathbf{O}}_{bp2} - \vec{\mathbf{O}}_{bp1})\cdot\widehat{\mathbf{h}} \tag{S39}$$

The algorithm then determines the angle $\phi$, which is the angle between $\widehat{\mathbf{H}}_1$ and $\widehat{\mathbf{y}}'_{hs1}$ (or between $\widehat{\mathbf{H}}_2$ and $\widehat{\mathbf{y}}'_{hs2}$) given the directionality of $\widehat{\mathbf{z}}_{hs}$ (S40). Using this angle, we can calculate Tip $\theta$ (see Figure S10e) (S41) and Inclination $\eta$ (see Figure S10d) (S42).

$$\phi = (\cos^{-1}(\widehat{\mathbf{H}}_1\cdot\widehat{\mathbf{y}}'_{hs1})\frac{180}{\pi})(\operatorname{sgn}((\widehat{\mathbf{y}}'_{hs1}\times\widehat{\mathbf{H}}_1)\cdot\widehat{\mathbf{z}}_{hs})) \tag{S40}$$

$$\theta = \Psi\cos(\phi) \tag{S41}$$

$$\eta = \Psi\sin(\phi) \tag{S42}$$

Next, similar to how it is done in the X3DNA/DSSR algorithms, it is necessary to determine the position of the origin of the local helical frame for any base pair in the step using Chasles' theorem (S43, S44, S45, S46). We always use base pair 1, but it is important to keep in mind that, due to the symmetric definition, using base pair 2 will give similar results.

$$\overrightarrow{\mathbf{AB}} = (\vec{\mathbf{O}}_{bp2} - \vec{\mathbf{O}}_{bp1}) - (d_z\cdot\widehat{\mathbf{h}}) \tag{S43}$$

$$\alpha = 90^\circ - (0.5\Omega) \tag{S44}$$

$$\widehat{\mathbf{AD}} = \frac{\mathbf{R}_h(\alpha)\cdot\overrightarrow{\mathbf{AB}}}{\|\overrightarrow{\mathbf{AB}}\|} \tag{S45}$$

$$AD_m = \frac{\|\overrightarrow{\mathbf{AB}}\|}{2\sin(\frac{\Omega}{2}\frac{\pi}{180})} \tag{S46}$$

Knowing $AD_m$ and the direction $\widehat{\mathbf{AD}}$, one can determine the positions of the origins of the local helical frames $\vec{\mathbf{O}}_{h1}$ (S47) and $\vec{\mathbf{O}}_{h2}$ (S48).

$$\vec{\mathbf{O}}_{h1} = \vec{\mathbf{O}}_{bp1} + (AD_m \cdot \widehat{\mathbf{AD}}) \tag{S47}$$

$$\vec{\mathbf{O}}_{h2} = \vec{\mathbf{O}}_{h1} + (d_z \cdot \widehat{\mathbf{h}}) \tag{S48}$$

The origin of a helical step is defined as the midpoint between the origins of the local helical frames (S49).

$$\vec{\mathbf{O}}_h = \frac{\vec{\mathbf{O}}_{h1} + \vec{\mathbf{O}}_{h2}}{2} \tag{S49}$$

The x-displacement $d_x$ (see Figure S10a) (S50) and the y-displacement $d_y$ (see Figure S10b) (S51) are defined as projections of the vector going from $\vec{\mathbf{O}}_{h1}$ to $\vec{\mathbf{O}}_{bp1}$ onto the corresponding components of the transformed basis triad $\mathbf{T}'_{hs1}$.

$$d_x = (\vec{\mathbf{O}}_{bp1} - \vec{\mathbf{O}}_{h1}) \cdot \widehat{\mathbf{x}}'_{hs1} \tag{S50}$$

$$d_y = (\vec{\mathbf{O}}_{bp1} - \vec{\mathbf{O}}_{h1}) \cdot \widehat{\mathbf{y}}'_{hs1} \tag{S51}$$

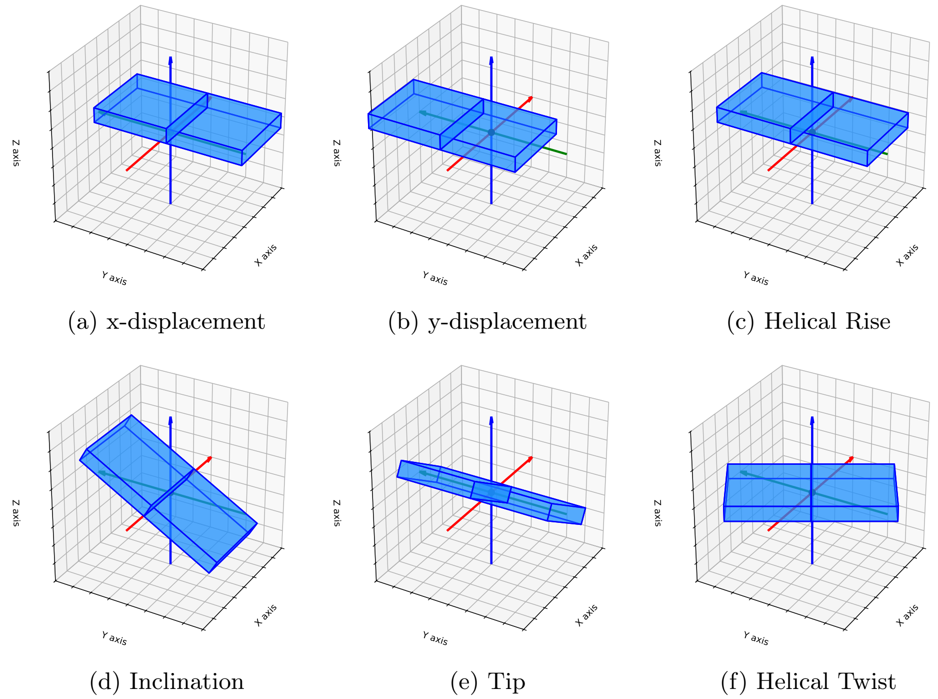


(a) x-displacement (b) y-displacement (c) Helical Rise

(d) Inclination (e) Tip (f) Helical Twist

Figure S10: Six rigid-body parameters of helical steps.

S.XXIV The DSSNA algorithm can distinguish A-like and B-like helical forms of nucleic acid dinucleotide steps by analyzing the relative positions of phosphorus atoms (see S.XXV). To determine the helix type in DSSNA, a straightforward criterion based on the $p_{dz}$ and $p_{hz}$ projection values is used[1].

1. If $p_{dz} \geq 0.15$ nm, then the dinucleotide step is assigned an A-like helical form;
2. If $p_{dz} \leq 0.05$ nm and $p_{hz} \leq 0.4$ nm, then the dinucleotide step is assigned a B-like helical form.

3. If a conformation does not satisfy either of these criteria, it is not assigned to the A-like or B-like helical forms.

S.XXV To determine the A-like and B-like helical forms of a dinucleotide step, we analyze the projection of the phosphate vector onto the dinucleotide frame and the helical frame. The phosphate vector is defined as the vector between the phosphorus atoms of the two nucleotides that form the dinucleotide step.

The algorithm calculates the projections $p_{dx}$, $p_{dy}$, and $p_{dz}$ of the vector $\vec{\mathbf{p}}_d$ (S52) and the projections $p_{hx}$, $p_{hy}$, and $p_{hz}$ of the vector $\vec{\mathbf{p}}_h$ (S53).

$$\vec{\mathbf{p}}_d = (\vec{\mathbf{p}}_1 - \vec{\mathbf{p}}_2) \cdot \mathbf{T}_{ds} \tag{S52}$$

$$\vec{\mathbf{p}}_h = (\vec{\mathbf{p}}_1 - \vec{\mathbf{p}}_2) \cdot \mathbf{T}_{hs} \tag{S53}$$

S.XXVI A helix is a sequential chain of at least two consecutive base pairs of any type. The base pairs do not necessarily have to be connected to one another through the sugar-phosphate backbone, but each neighboring pair of base pairs must be linked by at least one base-stacking interaction (see S.XXVII)[9].

The procedure used by DSSNA to traverse the structure and identify helices is as follows:

1. An arbitrary base pair is selected, typically the base pair located closest to the first nucleotide in the structure.
2. All other base pairs that are connected to the selected base pair through base-stacking interactions are identified. A connected base pair is excluded if it already belongs to another helix, is a pseudo-pair, or is adjacent to a base pair that has already been traversed.
3. The search proceeds toward the base pair with the greatest similarity to Watson–Crick geometry (see S.XXVIII). This base pair is added to the helix, and the procedure is repeated until no eligible base pairs remain.

It should be noted that helix types are not assigned to the secondary-structure motifs themselves. Instead, they are determined for the dinucleotide steps forming the structure.

DSSNA also provides an option for constructing a helix vector (see Figure S11). The vector is obtained by performing a principal component analysis of the origins of the base pairs forming the helix. The helix origin is defined as the centroid of all component base-pair origins. The direction of the helix vector is chosen to coincide with the vector extending from the origin of the first base pair to the origin of the second base pair in the helix.

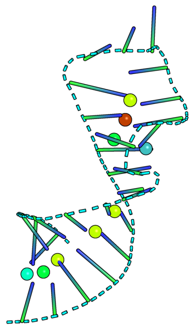

(a)

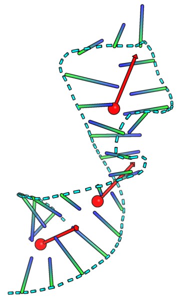

(b)

Figure S11: (S11a) Base pair origins that form different helices. (S11b) After base pair origins have been grouped together by base-stacking interactions, helical vectors are formed using principal component analysis.

S.XXVII Two distinct base pairs are considered to be involved in a base-pair stacking interaction if at least one nucleobase from the first base pair forms a base-stacking interaction with a nucleobase from the second base pair (see S.XII).

S.XXVIII During helix construction, the DSSNA algorithm may encounter a situation in which two adjacent base pairs are both eligible for inclusion in the helix. In this case, it is necessary to determine toward which base pair the helix should be extended.

Following the approach used in X3DNA/DSSR, the helix is extended toward the base pair that is geometrically closer to an idealized Watson–Crick base pair. To make this determination, we use a parameter based on the angular deviations of the nucleotide base-frame axes from their idealized orientations.

In an ideal Watson–Crick base pair, the $\widehat{\mathbf{x}}_1$ and $\widehat{\mathbf{x}}_2$ axes of the two nucleotides are parallel; that is, the angle between them is close to 0°. The $\widehat{\mathbf{y}}_1$ and $\widehat{\mathbf{y}}_2$ axes, as well as the $\widehat{\mathbf{z}}_1$ and $\widehat{\mathbf{z}}_2$ axes, are antiparallel; that is, the corresponding angles are close to 180°. Accordingly, we calculate the angular deviations $\alpha$ (S54), $\beta$ (S55), and $\gamma$ (S56) between the actual and idealized axis orientations. These values are then used to calculate the dispersion $D$ (S57). The helix is therefore preferentially extended toward the base pair with the smallest value of $D$, corresponding to the greatest geometric similarity to an ideal Watson–Crick base pair.

$$\alpha = 0 - \cos^{-1}(\widehat{\mathbf{x}}_1 \cdot \widehat{\mathbf{x}}_2)\frac{180}{\pi} \tag{S54}$$

$$\beta = 180 - \cos^{-1}(\widehat{\mathbf{y}}_1 \cdot \widehat{\mathbf{y}}_2)\frac{180}{\pi} \tag{S55}$$

$$\gamma = 180 - \cos^{-1}(\widehat{\mathbf{z}}_1 \cdot \widehat{\mathbf{z}}_2)\frac{180}{\pi} \tag{S56}$$

$$D = \sqrt{\frac{\alpha^2 + \beta^2 + \gamma^2}{3}} \tag{S57}$$

S.XXIX A stem is a segment of a helix with a continuous backbone, that is, without breaks in the sugar-phosphate backbone, and consisting exclusively of canonical base pairs[9]. By definition, a stem must contain at least two base pairs.

The presence of multiple stems within a single helix indicates a potential coaxial-stacking arrangement. This interpretation is valid when the terminal base pairs of neighboring stems are connected by base-stacking interactions and the corresponding helical axes are approximately collinear.

S.XXX An isolated canonical base pair is a canonical base pair that does not belong to a stem, which by definition consists of at least two base pairs. Isolated base pairs are of particular interest because they may serve as anchoring points for tertiary-structure interactions and help to "isolate" loops and pseudoknots[9].

S.XXXI An atom-base capping interaction is a stacking interaction in which an atom from a phosphate, sugar, or water molecule is positioned above the ring of a nucleobase, thereby capping the base and, in some cases, the end of a helix. The phosphate group is most commonly represented by its exocyclic OP2 atom. In addition to phosphate atoms, oxygen atoms from sugar moieties and water molecules can also serve as stacking atoms[9].

The atom-base capping interaction is considered to be formed if the following criteria are met for at least one geometric center of the aromatic ring of the base:

1. The capping atom, which is determined by selecting atoms of specific chemical elements, is no further than 0.55 nm (the value can be varied by the user) from the base.

2. The vertical separation (see S.X) of the capping atom relative to the geometric center of the aromatic ring of the base exceeds 0.35 nm (the value can be varied by the user).
3. The angle between the vector extending from the geometric center of the base to the position of the capping atom in space and the base normal exceeds 25° (the value can be varied by the user).

DSSNA searches for oxygen atom caps by default, but the user has the option to select the element type to search for. Since base capping interactions are searched within nucleic acids, detection of base capping by water oxygen atoms is currently not supported.

S.XXXII Single-stranded fragments are continuous sequences of nucleotides that do not belong to any helix, stem, isolated base pair, or loop[9].

S.XXXIII Multiplets (see Figure S12) are tertiary nucleic motifs. They represent higher-order coplanar nucleotide associations[9]. In simple terms, they are several nucleotides linked by a hydrogen bond network. The most important multiplet characteristic is its size — the number of nucleotides comprising the multiplet. The size of a multiplet cannot be less than 3, but technically there is no upper limit.

In DSSNA, multiplets are identified using hydrogen bond network analysis. The first nucleotide from the sequence is taken, other nucleotides linked to it by hydrogen bonds (if any) are identified, and they are added to a single set. The algorithm is then repeated for other nucleotides linked to the second nucleotide. The algorithm is repeated until there are no more nucleotides linked by other hydrogen bonds, and all linked nucleotides are added to a single set (if a nucleotide is isolated, the set will consist only of that nucleotide). If the size of the final set reaches the minimum (3 nucleotides), the set will be reflected in the final output as a multiplet.

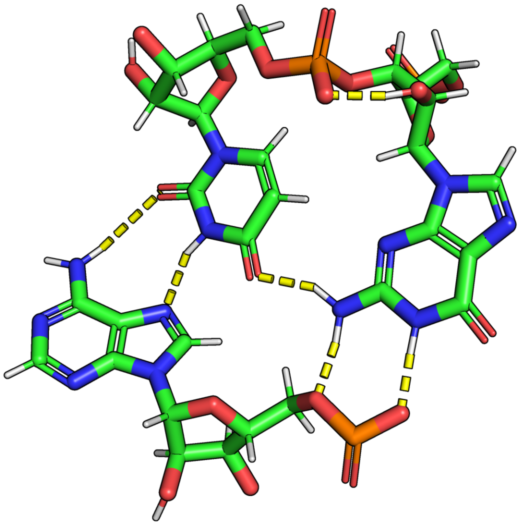

Figure S12: Visualization of a multiplet of size 3 (with hydrogen bond network).

S.XXXIV A minor motif (see Figure S13) is a tertiary nucleic (mostly RNA) motif. It is a special case of a base triplet (multiplet of fixed size 3) that represents a canonical base pair linked by one or more hydrogen bonds to a nucleotide whose base (the "interacting base") is of a selected type. Most often, this base is adenine (the motif is then called an "A-minor motif")[11]. Accordingly, DSSNA is configured by default to detect A-minor motifs. The user can also configure the algorithm to detect analogous motifs involving other nucleobase types. In such cases, the detected structures are more generally referred to as minor motifs or A-minor-like motifs.

DSSNA implements three operational classes of minor-groove base-triple interactions: Types I, II, and X. These classes are defined according to the hydrogen-bonding pattern and the relative orientation of the interacting base and the receptor base pair. They should not be regarded as a direct replacement for the classical Nissen classification[12], which is based primarily on the positions of the O2' and N3 atoms of adenine relative to the receptor minor groove. While DSSNA takes these atoms into account when searching for hydrogen bonds in A-minor motifs, its definition of minor motifs is deliberately made broader.

1. Type I (see Figure S13a) — The interacting base presents at least one minor-groove-edge atom that forms at least one hydrogen bond with each of the two canonical bases of the receptor base pair.
2. Type II (see Figure S13b) — The interacting base presents at least one minor-groove-edge atom that forms at least one hydrogen bond with one of the canonical bases of the receptor base pair.
3. Type X (see Figure S13c) — The interacting base presents at least one major-groove-edge atom that forms at least one hydrogen bond with one of the canonical bases of the receptor base pair through the minor-groove side of the pair.

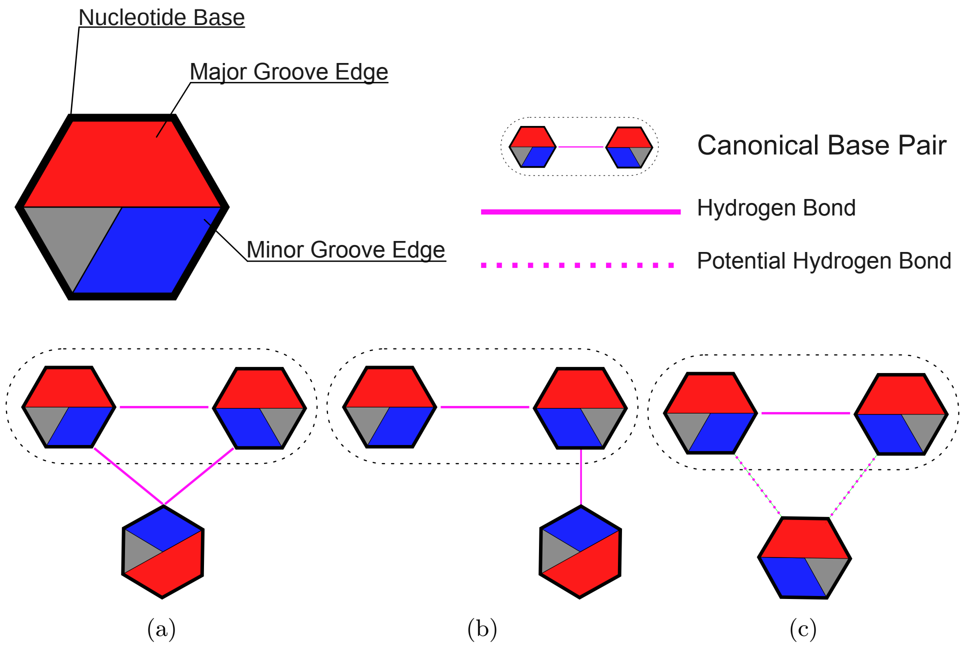


Figure S13: Minor motif diagram. (S13a) Minor motif Type I. (S13b) Minor motif Type II. (S13c) Minor motif Type X.

S.XXXV A ladder (see Figure S14) is a secondary nucleic motif. It consists of two consecutive nucleotide chains on one (see Figure S14a) or two (see Figure S14b) nucleic acid strands, with at least one non-overlapping base pair between them. Ladders can sometimes conflict with each other, forming nodes between them. Ladders have upstream (5'-end) and downstream (3'-end) regions[13].

Ladders are characterized by parameters such as the number of hydrogen bonds in the ladder, length (the number of base pairs in the ladder), range (the difference between the highest upstream position (closest to the 5' end) and the lowest downstream position (closest to the 3' end)) and gain (the "gain" of accepting base pairs in this ladder, defined as the length minus the sum of all the lengths of regions that conflict with this ladder).

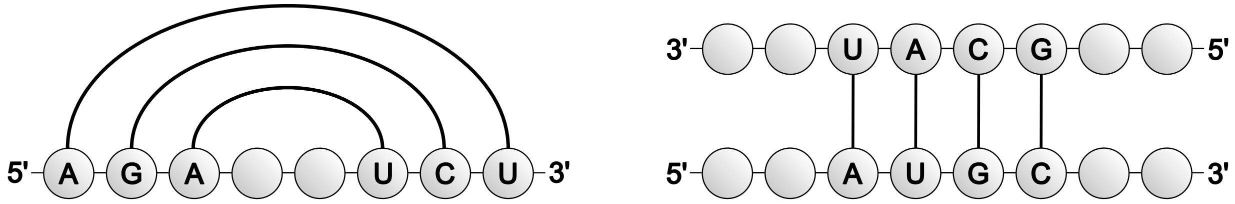


(a) (b)

Figure S14: Ladder motif diagram. (S14a) Ladder in a single chain. (S14b) Ladder in a double chain.

S.XXXVI A G-tetrad (also known as a G-quartet; see Figure S15) is a planar, approximately square arrangement of four guanine bases connected by a cyclic network of Hoogsteen-type hydrogen bonds. It is a fundamental structural unit of a G-quadruplex. By definition, the G-tetrad is a multiplet of size 4. Monovalent cations are usually located in the central channel of the tetrad and provide additional electrostatic stabilization[14].

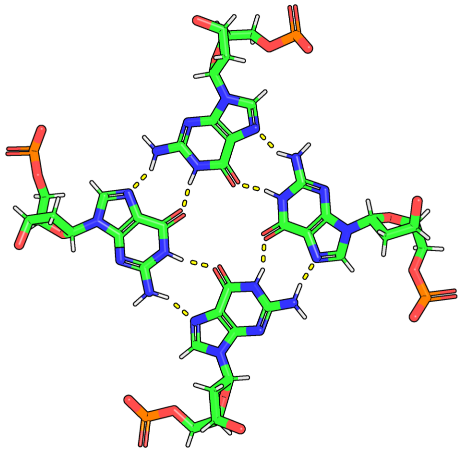

Figure S15: Visualization of a G-tetrad (a G+G motif that forms G-quadruplexes) with its hydrogen-bond network.

S.XXXVII A loop (see Figure S16) is an element of the secondary structure of nucleic acids. Loops are closely associated with the secondary structure, as they are delimited by canonical base pairs held together by hydrogen bonds and base-stacking interactions. Understanding loop structure is essential for predicting nucleotide interactions in a dynamic profile, especially for RNA.

Loops come in different types (see S.XXXVIII), some of which may exhibit symmetry properties (see S.XXXIX). Loops have one or more internal parts (designated as I in the output file) and spacer nucleotides (delimiters, designated as D in the output file) that delimit the internal parts of the loop. Delimiters are formed from canonical base pairs that form stems or isolated base pairs.

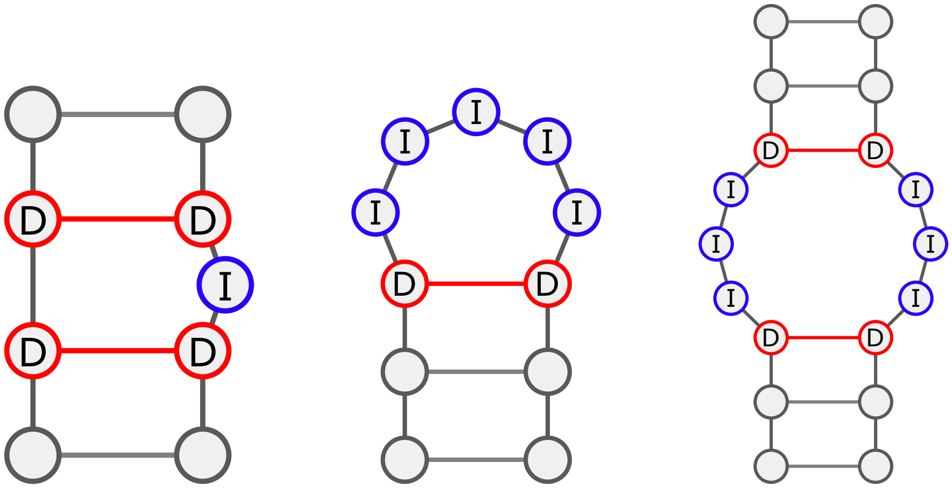


(a) (b) (c)

Figure S16: Loop formation logic diagram. I stands for “internal nucleotide”. D stands for “delimiter nucleotide”. (S16a) Bulge. (S16b) Hairpin loop. (S16c) Internal loop. The image was generated using VARNA[15].

S.XXXVIII The secondary-structure loop motifs defined in DSSNA (see Figure S17) are classified as follows:

1. Undefined — A technical classification for loops that, for some reason, could not be assigned a specific type.
2. Bulge — Loops that have only one set of internal nucleotides and whose delimiter consists of more than 2 nucleotides.
3. Internal loop — Loops that have exactly 2 sets of internal nucleotides with exactly 2 delimiter nucleotides.
4. Hairpin loop — Loops that have exactly one set of internal nucleotides with exactly 2 delimiter nucleotides.
5. N-way junction — Loops that have more than 2 sets of internal nucleotides. They are sometimes also denoted using the nucleotide count of the internal parts.

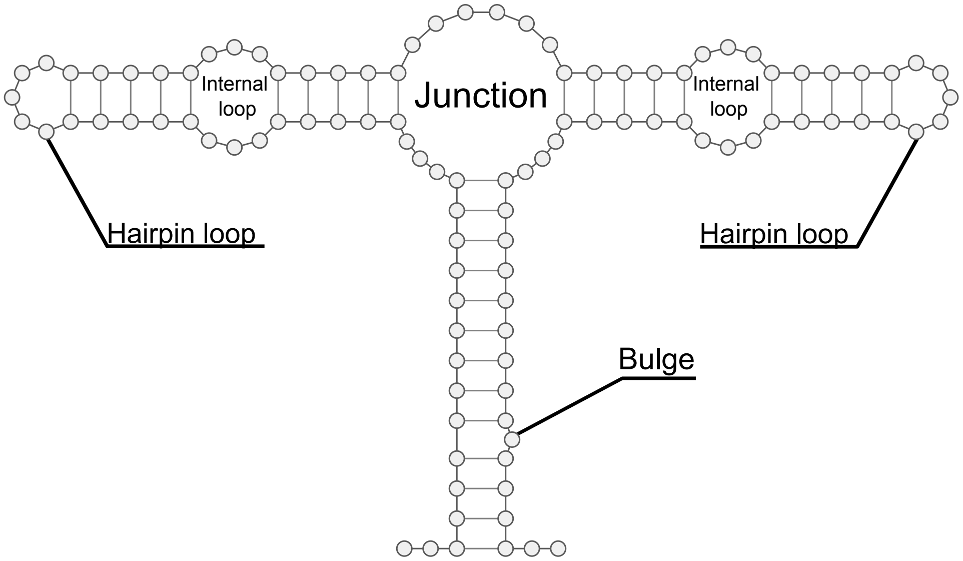


Figure S17: Diagram of the different types of loops that can be found in DSSNA. The image was generated using VARNA[15].

Only secondary motifs that form loops are described above. The kissing loops motif (see S.XL) is a tertiary motif and is calculated separately.

S.XXXIX DSSNA defines the following loop symmetry types:

1. Undefined — Loops with only one set of internal nucleotides cannot have any defined symmetry type.
2. Symmetrical — Loops with the same number of nucleotides in their sets of internal nucleotides.
3. Asymmetrical — Loops with different numbers of nucleotides in their sets of internal nucleotides.

S.XL A kissing loop (see Figure S18) is a tertiary nucleic motif. It is a type of intermolecular or intramolecular interaction in nucleic acids, in which two single-stranded nucleic acid molecules (or different sections of a single strand) recognize each other and join through the formation of hydrogen bonds between complementary nucleotides (thus forming canonical base pairs) located at the tips of two hairpins. It predominantly forms in RNA. Kissing loops often play an important role in RNA folding and stabilizing its structure.

The interaction occurs according to the “handshake” principle: first, several initial nucleotides in the loop are paired. This ensures rapid and reversible recognition without the large energy expenditure required to unwind stable helices.

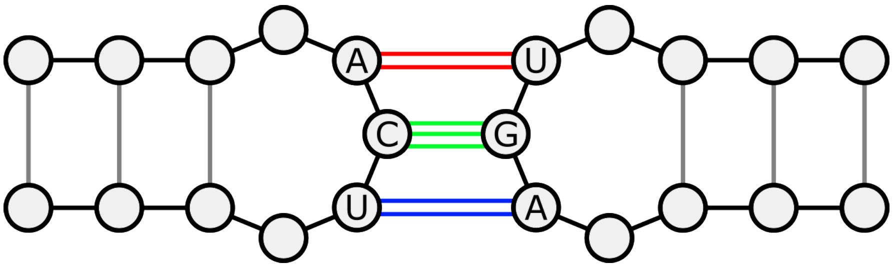


Figure S18: Kissing loop motif schematic representation. The image was generated using VARNA[15].

DSSNA defines kissing loops as canonical base pairs (or, optionally, any non-pseudo-pairs) between hairpin loops.

S.XLI Pseudoknots are atypical elements of the secondary structure of nucleic acids (primarily RNA due to their high mobility), representing spatial overlaps of base pairs. If we decompose a nucleic acid linearly and draw the base pairs as lines between nucleotides, we obtain a ladder map (see S.XXXV). When these ladders overlap, they form pseudoknots (see Figure S19).

Identifying pseudoknots is essential for correctly predicting and understanding the secondary/tertiary structure of RNA, which regulates functions such as translation, splicing, ribozyme activity, and frameshifting in viruses or genes.

Often, due to limitations in algorithms or programs, it is not possible to work with structures containing pseudoknots. Their presence can radically alter the behavior of software algorithms (including DSSNA), and their removal may be necessary for comparing structures. Therefore, we have implemented several algorithms that allow us to identify pseudoknots and, if desired, remove them.

Identification begins with constructing a ladder map formed by base pairs for the selected structure/trajectory. Next, the presence of nodes (conflicts) between these ladders is checked. The nodes are evaluated using various metrics to determine which node should be retained. The evaluation algorithms for most methods are implemented according to the methods described in Smit *et al.* (2008)[13].

We have developed an analog of an optimization algorithm that analyzes all potential sets of pseudoknot configurations and selects the best one using a scoring function that maximizes the number of hydrogen bonds in base pairs. This is the default algorithm. All base pairs that form knots will be removed from the structure and will not be considered in further analysis. Pseudoknot removal is optional and can be omitted at the user's discretion.

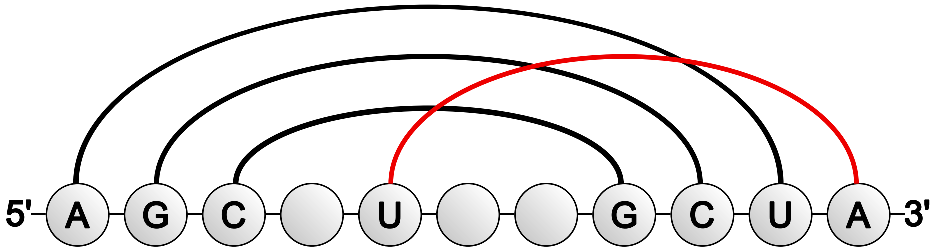


Figure S19: Schematic representation of a pseudoknot in a ladder.

S.XLII Dot-bracket notation (DBN) is a compact text format used to represent nucleic acid secondary structures. DBN uses dots to represent nucleotides that do not form base pairs and brackets to represent nucleotides that are linked by base pairs. When the number of brackets runs out, letters of the English alphabet are used (uppercase at the beginning, lowercase at the end).

In DSSNA, the dot-bracket notation is generated using a standardized one-letter representation of nucleotides. Modified nucleotides are automatically converted to their closest canonical counterparts in the DBN output. This ensures compatibility with standard secondary-structure visualization tools and established DBN conventions, while the original residue names from the topology are retained in the primary output files for reference.

S.XLIII An i-motif is a special type of secondary structure that occurs in cytosine-rich regions of nucleic acids (usually DNA)[16]. The i-motif is classically defined as a four-stranded intercalated structure stabilized by hemiprotonated cytosine-cytosine base pairs $C{+}C^{+}$.

The building block of the i-motif is generally considered to be a hemiprotonated cytosine-cytosine base pair $C{+}C^{+}$ (see Figure S20a). However, the protonation state of cytosines is sometimes unavailable in coordinate-based structural analyses. Therefore, DSSNA can identify i-motif-like geometries based on the spatial arrangement and pairing pattern of cytosine residues, without requiring explicit protonation-state information (see Figure S20b).

DSSNA implements a dedicated procedure for identifying i-motif (and i-motif-like) structures, based on an approach analogous to the helix-detection algorithm. The procedure first identifies candidate $C{+}C^{+}$

base pairs and then searches for groups of such pairs connected by base-stacking interactions. These groups are treated as structural i-motif blocks according to the criteria defined for the corresponding motif class.

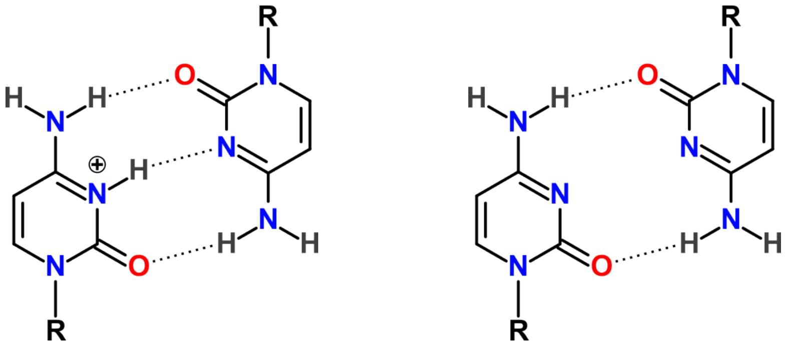

(a) (b)

Figure S20: (S20a) Standard definition of i-motif building blocks where one cytosine is partially protonated (C+C$^+$ pair). (S20b) Extended geometric definition of i-motif-like building blocks without protonated cytosine (C+C pair).

S.XLIV A splayed apart conformation is the spatial arrangement of two consecutive nucleotides that are spaced far apart. Splayed conformations are calculated by analyzing the positions of the P atom connecting the nucleotides, the O3' atoms, and the nucleobase origins.

To calculate splayed conformations, we find the vector between the origins of the two nucleotides, $\overrightarrow{\mathbf{O_1O_2}}$, and the vector $\overrightarrow{\mathbf{PO3'}}$, such that $\|\overrightarrow{\mathbf{PO3'}}\| \leq 0.159 \cdot 1.25$ nm (the average O3'–P bond length with the same 25% variability allowance that is used for breaks; see S.XLVII).

The output file specifies the splayed angle $\alpha$ (S58), distance $d$ (S59), and ratio $r$ (S60).

$$\alpha = \cos^{-1}(\widehat{\mathbf{PO_1}} \cdot \widehat{\mathbf{PO_2}})\frac{180}{\pi} \tag{S58}$$

$$d = \|\overrightarrow{\mathbf{O_1O_2}}\| \tag{S59}$$

$$r = \frac{\|\overrightarrow{\mathbf{O_1O_2}}\|}{\|\overrightarrow{\mathbf{PO_1}}\| + \|\overrightarrow{\mathbf{PO_2}}\|} \tag{S60}$$

S.XLV The backbone is the structural framework that holds the nucleic structure together, consisting of alternating sugar (ribose or deoxyribose) and phosphoric acid residues. It connects nucleotides into a single chain. The backbone is described by many parameters, such as turn (see S.XLVI), break (see S.XLVII), dihedral angles (see S.XLVIII), virtual dihedral angles (see S.XLIX), and sugar conformation (see S.L).

S.XLVI Turn is a backbone parameter determined by analyzing the angle between successive C1'–C1' atoms. For example, for nucleotide i, two vectors, $\overrightarrow{\mathbf{C1'_{i-1}C1'_{i}}}$ and $\overrightarrow{\mathbf{C1'_{i}C1'_{i+1}}}$, are calculated, and the angle between them is then determined. If the angle exceeds 90°, the sugar-phosphate backbone of nucleotide i is assigned a secondary turn motif. Even if nucleotides in the chain have a break (before and/or after), their turn can still be analyzed.

S.XLVII The break between two nucleotides is determined by assessing the distance between the O3' and P atoms. For each nucleotide pair, two pairs of lengths between these atoms are checked (if present in the structure). If the length between the atoms in at least one pair is less than the established cutoff, then the break between the nucleotides is considered to be absent. The average length of the O3'–P covalent bond, 0.159 nm[17], was chosen as the cutoff, with an additional allowance for variability (25%) so that values slightly exceeding the average are not discarded.

S.XLVIII Dihedral (torsion) angles are mathematical measurements that describe the 3D shape, orientation, and flexibility of the DNA/RNA backbone, sugar rings, and base pairs (see Figure S21).

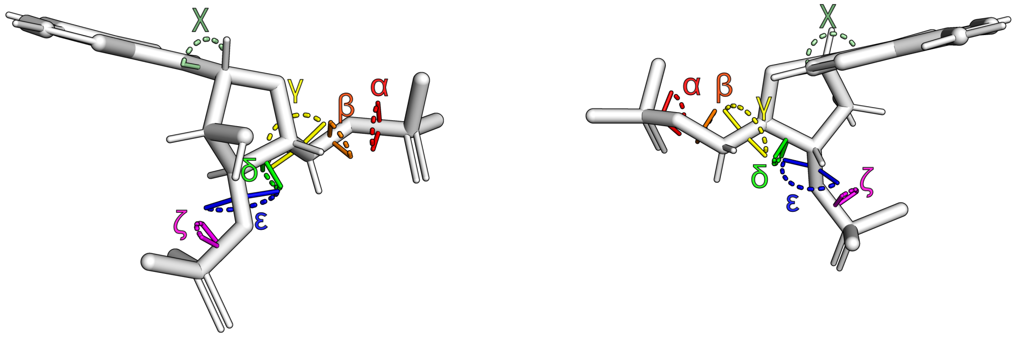


Figure S21: Diagram of the seven standard torsion angles in the nucleic acid backbone.

There are seven standard torsion angles: $\alpha$, $\beta$, $\gamma$, $\delta$, $\varepsilon$, $\zeta$ and $\chi$[18]. They differ from each other in the atoms that are used to define them (see Table S4).

Table S4: Atoms used in dihedral (torsion) angle calculation for nucleotide $i$.

| Dihedral angle | Atom A | Atom B | Atom C | Atom D |
|---|---|---|---|---|
| Alpha ($\alpha$) | $O3'_{i-1}$ | $P_i$ | $O5'_i$ | $C5'_i$ |
| Beta ($\beta$) | $P_i$ | $O5'_i$ | $C5'_i$ | $C4'_i$ |
| Gamma ($\gamma$) | $O5'_i$ | $C5'_i$ | $C4'_i$ | $C3'_i$ |
| Delta ($\delta$) | $C5'_i$ | $C4'_i$ | $C3'_i$ | $O3'_i$ |
| Epsilon ($\varepsilon$) | $C4'_i$ | $C3'_i$ | $O3'_i$ | $P_{i+1}$ |
| Zeta ($\zeta$) | $C3'_i$ | $O3'_i$ | $P_{i+1}$ | $O5'_{i+1}$ |
| Chi ($\chi$) in purines | $O4'_i$ | $C1'_i$ | $N9_i$ | $C4_i$ |
| Chi ($\chi$) in pyrimidines | $O4'_i$ | $C1'_i$ | $N1_i$ | $C2_i$ |

Additionally, the difference between the dihedral angles $\varepsilon$ and $\zeta$ (E-Z) is calculated. If the difference exceeds 180°, 360° is subtracted from it. If the difference is less than −180°, 360° is added to it. If the final value of E-Z $\in [-160°; 20°]$, then the backbone is assigned the $B_I$ type. If the resulting E-Z value is $\in (20°; 200°]$, then the backbone is assigned the $B_{II}$ type[18].

The base conformation is assigned depending on the value of the angle $\chi$. If the angle $\chi \in [-180°; -50°] \bigcup [170°; 180°]$, then the base is assigned the *anti* conformational type. If $\chi \in [50°; 90°]$, then the base is assigned the *syn* conformational type[9].

The nucleotide parameters $ssZ_p$, $D_p$, and *splay* are determined if possible. $ssZ_p$ — stands for "single-stranded" phosphorus z-axis projection — the projection of the vector $\overrightarrow{\mathbf{OP}}$, running from the nucleotide origin $\vec{\mathbf{O}}$ to the phosphorus atom P of the next nucleotide in the chain, onto the nucleotide normal $\widehat{\mathbf{n}}$ (S61).

$$ssZ_p = \widehat{\mathbf{n}} \cdot \overrightarrow{\mathbf{OP}} \tag{S61}$$

$D_p$ is the perpendicular distance from the 3' phosphorus (P) atom to the glycosidic bond (the bond connecting the sugar to the base) of its own nucleotide. To calculate it, two vectors must be determined:

the $\overrightarrow{\mathbf{C1'P}}$ vector, extending from the C1' atom of the nucleotide to the phosphorus P atom of the next nucleotide, and the (normalized) $\widehat{\mathbf{C1'N}}$ vector, extending from the C1' atom of the nucleotide to the nitrogen atom that forms the glycosidic bond (N9 for purines, N1 for pyrimidines). Knowing these two vectors, the exact value of the $\overrightarrow{\mathbf{D_p}}$ projection can be determined, and the $D_p$ value will be its norm (S62).

If the calculated value of $D_p > 0.29\ nm$, then the sugar-phosphate backbone is assigned the type *C3'-endo*; otherwise, it is assigned the type *C2'-endo*.

$$\begin{aligned} \overrightarrow{\mathbf{D_p}} &= \overrightarrow{\mathbf{C1'P}} - ((\overrightarrow{\mathbf{C1'P}} \cdot \widehat{\mathbf{C1'N}}) \cdot \widehat{\mathbf{C1'N}}) \\ D_p &= \|\overrightarrow{\mathbf{D_p}}\| \end{aligned} \tag{S62}$$

*splay* (or *splay angle*) is the angle between the two origins $\overrightarrow{\mathbf{O_1}}$ and $\overrightarrow{\mathbf{O_2}}$ of two adjacent backbone nucleotides and the phosphorus atom P connecting them. This metric is used to quantify how sharply a nucleic acid backbone "opens up" or bends between adjacent nucleotides (S63).

$$splay = \cos^{-1}(\widehat{\mathbf{PO_1}} \cdot \widehat{\mathbf{PO_2}})\frac{180}{\pi} \tag{S63}$$

S.XLIX Virtual dihedral (torsion) angles are simplified mathematical measurements used to describe the complex, twisting backbone geometry of nucleic acids (see Table S5).

Table S5: Atoms used in virtual dihedral (torsion) angle calculation for nucleotide $i$.

| Dihedral angle | Atom/Origin A | Atom/Origin B | Atom/Origin C | Atom/Origin D |
|---|---|---|---|---|
| Eta ($\eta$) | C4'$_{i-1}$ | P$_i$ | C4'$_i$ | P$_{i+1}$ |
| Theta ($\theta$) | P$_i$ | C4'$_i$ | P$_{i+1}$ | C4'$_{i+1}$ |
| Eta' ($\eta$') | C1'$_{i-1}$ | P$_i$ | C1'$_i$ | P$_{i+1}$ |
| Theta' ($\theta$') | P$_i$ | C1'$_i$ | P$_{i+1}$ | C1'$_{i+1}$ |
| Eta" ($\eta''$) | O$_{i-1}$ | P$_i$ | O'$_i$ | P$_{i+1}$ |
| Theta" ($\theta''$) | P$_i$ | O$_i$ | P$_{i+1}$ | O$_{i+1}$ |

S.L To accurately determine the conformation of the sugar-phosphate backbone of a nucleic acid, 5 torsion angles are used: $v_0$, $v_1$, $v_2$, $v_3$ and $v_4$ (see Table S6).

Table S6: Atoms used in sugar-ring torsion angle calculation for nucleotide $i$.

| Dihedral angle | Atom A | Atom B | Atom C | Atom D |
|---|---|---|---|---|
| $v_0$ | C4'$_i$ | O4'$_i$ | C1'$_i$ | C2'$_i$ |
| $v_1$ | O4'$_i$ | C1'$_i$ | C2'$_i$ | C3'$_i$ |
| $v_2$ | C1'$_i$ | C2'$_i$ | C3'$_i$ | C4'$_i$ |
| $v_3$ | C2'$_i$ | C3'$_i$ | C4'$_i$ | O4'$_i$ |
| $v_4$ | C3'$_i$ | C4'$_i$ | O4'$_i$ | C1'$_i$ |

Using the five torsion angles listed above, the amplitude of puckering of a nucleic acid's sugar ring (ribose or deoxyribose) $t_m$ and sugar pucker phase angle $P$ are calculated. The DSSNA algorithm implements two methods for calculating these parameters: one reproduces the calculation used in X3DNA[1] (S64, S65)

$$\begin{aligned} a &= v_4 + v_1 - v_3 - v_0 \\ b &= 2v_2 \sin(\frac{\pi}{5}) + \sin(\frac{\pi}{2.5}) \\ P &= \text{atan2}(a, b)\frac{180}{\pi} \end{aligned} \tag{S64}$$

$$t_m = \frac{v_2}{\cos(P)} \tag{S65}$$

and the other used in Curves+[19] (S67, S66).

$$\begin{aligned} a &= 0.4(v_2 + v_3\cos(0.8\pi) + v_4\cos(1.6\pi) + v_0\cos(2.4\pi) + v_1\cos(3.2\pi)) \\ b &= 0.4(v_3\sin(0.8\pi) + v_4\sin(1.6\pi) + v_0\sin(2.4\pi) + v_1\sin(3.2\pi)) \\ t_m &= \sqrt{a^2 + b^2} \end{aligned} \tag{S66}$$

$$P = \text{atan2}(\frac{a}{t_m}, \frac{b}{t_m})\frac{180}{\pi} \tag{S67}$$

If $P < 0$, then 360° is added to the value of $P$. Therefore, in both cases, $P \in [0°; 360°)$. Depending on the value of the phase angle $P$, it is assigned one of ten sugar pucker types (see Table S7).

Table S7: Ranges of the sugar pucker phase angle $P$ for all conformations of ribose or deoxyribose[18].

| **Angle range** | **Name** |
|---|---|
| $[0°; 36°)$ | C3'-endo |
| $[36°; 72°)$ | C4'-exo |
| $[72°; 108°)$ | O4'-endo |
| $[108°; 144°)$ | C1'-exo |
| $[144°; 180°)$ | C2'-endo |
| $[180°; 216°)$ | C3'-exo |
| $[216°; 252°)$ | C4'-endo |
| $[252°; 288°)$ | O4'-exo |
| $[288°; 324°)$ | C1'-endo |
| $[324°; 360°)$ | C2'-exo |

S.LI To calculate the dihedral angle $\theta$, four coordinates A, B, C, and D are required, which are usually nucleic acid atoms (but sometimes nucleotide origins as well). Vectors $\overrightarrow{\mathbf{BA}}$, $\overrightarrow{\mathbf{CB}}$, and $\overrightarrow{\mathbf{CD}}$ are determined. The normal $\vec{\mathbf{n}}_1$ (S68) is determined for the $ABC$ plane, and the normal $\vec{\mathbf{n}}_2$ (S69) is determined for the $BCD$ plane. Then, the components $x$ (S70) and $y$ (S71) are calculated, which are used to determine the dihedral angle $\theta$ (S72).

$$\vec{\mathbf{n}}_1 = \overrightarrow{\mathbf{CB}} \times \overrightarrow{\mathbf{BA}} \tag{S68}$$

$$\vec{\mathbf{n}}_2 = \overrightarrow{\mathbf{CB}} \times \overrightarrow{\mathbf{CD}} \tag{S69}$$

$$x = \frac{\vec{\mathbf{n}}_1 \cdot \vec{\mathbf{n}}_2}{\sqrt{\vec{\mathbf{n}}_2^2}} \tag{S70}$$

$$y = \frac{\vec{\mathbf{n}}_1 \cdot (\overrightarrow{\mathbf{CB}} \times \vec{\mathbf{n}}_2)}{\sqrt{(\overrightarrow{\mathbf{CB}} \times \vec{\mathbf{n}}_2)^2}} \tag{S71}$$

$$\theta = \text{atan2}(x, y)\frac{180}{\pi} \tag{S72}$$

By the definition above, the value of a dihedral angle in degrees can only lie in the range $\theta \in [-180°; 180°]$.

S.LII A ribose zipper is an RNA tertiary interaction between two distinct segments of the same RNA chain or between two different RNA chains. It is characterized by at least two consecutive nucleotides forming hydrogen bonds between their ribose 2'-hydroxyl groups, thereby bridging the two segments[20].

#### Implementation in terms of the software algorithms

In DSSNA, nucleotides are denoted using the following format: C.Nn, where C — optional chain identifier, N — nucleotide name from topology (or the most similar standard ribose/deoxyribose base if there is none), and n — nucleotide number in the structure. For example: "DNA_Chain_B.DA210" or "RNA_Chain_A.U67". Use the $-nochain$ option to remove chain identifiers from the nucleotide names.

When analyzing structures with DSSNA, particular attention should be paid to the treatment of the periodic boundary conditions (PBC). DSSNA relies extensively on distance calculations between pairs of points in space, with periodic boundary conditions taken into account. If the structure is associated with a simulation box that is too small, the calculated distances may be incorrect.

The $-o$ option is the standard (and primary) output option in DSSNA. It outputs information about nucleotide types and the spatial arrangements of their bases. In particular, it outputs information about nucleoframes (see S.I), namely their origin coordinates (see S.III), the unit vectors that form the right-hand orthonormal basis of the base, and the positions of the geometric centers of aromatic rings (AC — **A**romatic ring **C**enter). The orthonormal basis is formed using the least-squares fit method in the nucleotide type determination algorithm (see S.IV). This option is useful if you are interested in the geometric arrangement of nucleotides and/or want to visualize them.

The $-hbo$ option allows the user to obtain information about hydrogen bond networks based on hydrogen bond length and angle cutoffs (see S.V). The partial-charge threshold for atoms to be considered acceptors is determined by the $-energy$ parameter. The hydrogen bond length cutoff (the length between donor and acceptor) is specified by the user using the $-hbdist$ parameter. The hydrogen bond angle cutoff (hydrogen-donor-acceptor) is specified by the user using the $-hbang$ parameter. The information returned by this option allows users to analyze in detail the hydrogen bonds formed in the structure of nucleic acids. Note that this list displays not only hydrogen bonds formed between nucleic acid bases, but also hydrogen bonds formed by atoms of the sugar-phosphate backbone. Hydrogen bond parameters (like stacking interaction parameters) play a key role in determining the secondary structures of nucleic acids and should be modified with caution. Note that the hydrogen bond definition criteria in DSSNA and X3DNA/DSSR differ, which will lead to differences in base pair formation.

DSSNA offers a number of options that generate information on base pairs present in a structure/trajectory (see S.VIII). They all output the base pairs found in frames, but emphasize different types of information. All options share a common element: information about the two nucleotides that form the base pair, including abbreviations, relative orientation, type, and, if present, the number of hydrogen bonds in the $n + m'$ format, where $n$ is the number of hydrogen bonds between the bases and $m$ is the number of hydrogen bonds in the sugar-phosphate backbone. The $-bpso$ option outputs a list of base pairs with secondary structure identifiers, like helices (see S.XXVI), stems (see S.XXIX), and isolated pairs (see S.XXX). The $-bpfo$ option outputs a base pair reference (see S.VIII); it is useful if you want to display the pairs in a visualization utility or want to refine the absolute or relative positions of the base pairs of interest. The $-bpro$ option outputs the mutual geometric parameters of the base pair (see S.XXI). The $-bpang$ option specifies the degree of accuracy and controls the allowed deviation from optimal base pair geometry, which in turn affects the assignment of base pair types (see S.XV).

If you want to analyze the stability of base pairs or the total time it takes for them to form in a trajectory, use the $-bpoo$ and $-bpmo$ options. The $-bpoo$ option generates a map of the cumulative base-pair lifetimes over the MD trajectory. The $-bpmo$ option generates a map of the longest-lived base pairs in the trajectory, allowing you to evaluate their stability. The $-bpmmode$ option specifies which base pairs you want to use for this analysis: $-bpmmode$ *canon* (the default) uses only canonical base pairs, $-bpmmode$ *nonpseudo* — all non-pseudo base pairs, and $-bpmmode$ *all* — all base pairs. The $-mapmode$ option controls the data output format: $-mapmode$ *relation* (default) outputs map data as the ratio of lifetimes (total or only the longest-lived) to the total time of the MD trajectory (or its part); the $-mapmode$ *time* option outputs only the lifetime of structures.

The $-dso$ option allows you to obtain data regarding dinucleotide and helical steps (see S.XVIII). The output file will contain information about consecutive base pairs and the parameters of their rigid frames (see S.XXII, S.XXIII). This option also outputs information about the projection of the phosphate vector (see S.XXV) onto the rigid frames of the dinucleotide and helical steps as well as the helix type of the dinucleotide step (see S.XXIV). The rigid frame parameters presented include parameters related to the relative position of their rigid frames, as well as parameters related to the helix parameters of the dinucleotide step.

If you request an output containing a list of base pairs, false base pairs (pseudo-pairs) will not be included in it (by default). If you need to obtain lists that will include pseudo-pairs, you will need to specify the $-pseudo$ option. With that option on, pseudo-pairs will appear in the output and will have an asterisk (*) before their name.

The $-vo$ and $-bpvo$ options provide information about virtual bonds (see S.XVII) in nucleic acid structures. Option $-vo$ analyzes virtual bonds inside the nucleotide chain. Option $-bpvo$ examines virtual bonds between base pairs.

The $-ho$ option creates a file containing qualitative information about the helices detected in the molecular dynamics trajectory frames (see S.XXVI). The file contains information about the unique identifier of each helix, the number of base pairs it contains, and the number of stems (see S.XXIX) and isolated base pairs (see S.XXX) in each helix. This essentially reflects coaxial stacking (the relationship between helices and stems). Additionally, this file contains information about the reference points of each helix. This allows you to determine the center of each helix and its basis, which can be used for further visualization.

The $-bso$ option reports base-stacking interactions (see S.XII) detected in nucleic acid structures. The output file generated by this option contains a list of stacking interactions and their associated parameters. The algorithm implements two methods for identifying stacking interactions: estimation of the base-overlap area (option $-bssmode\ overlap$, default; see S.XIII) and analysis of geometric parameters (option $-bssmode\ geometry$; see S.XIV). In overlap-area mode, the minimum overlap ratio relative to the smaller base is controlled by the $-bsarea$ parameter. In geometric mode, the angular cutoff is specified by $-bsang2$, which defines the angle between the nucleotide normal and the normalized vector connecting the geometric centers of the two bases. In both modes, the distance cutoff between the geometric centers of the aromatic rings can be adjusted using $-bsdist$, and the angular cutoff between nucleobase normals can be adjusted using $-bsang1$. These cutoffs are applied not only when generating the stacking output file but also during stacking analysis in other parts of the program (for example, when searching for base pairs). By default, base-stacks within stems are omitted from the output file but can be included using the $-nohidebss$ option.

Similar to analyzing the stability of base pairs and the total time of their formation in a trajectory, you can analyze the stability of base-stacking interactions using the $-bsoo$ and $-bsmo$ options. The $-bsoo$ option generates a map of the total lifetime of stacking interactions throughout the entire MD trajectory (or its part). The $-bsmo$ option generates a map of the longest-lived stacking interactions in the trajectory. The $-mapmode$ option works similarly to the above. By default, base-stacks included in stems are not written to the output file, but can be added using the $-nohidebss$ option.

The $-bco$ option writes a file with the atom-base capping interactions within nucleic acids (see S.XXXI). The output file contains information about the atoms capping the nucleotides, their distance, vertical separation, and capping angle. By default, capping atoms are searched only for oxygen atoms, but you can specify elements of interest using the $-ba$ flag. Length, angle, and vertical separation cutoffs for capping atom searches are specified using the $-bcdist$, $-bcang$, and $-bcvs$ flags, respectively.

The $-lo$ option provides information about loops (see S.XXXVII). The module provides information on the total loop size, their symmetry, secondary structure, and the nucleotides that make up the loop. The maximum search depth is specified by the $-ld$ option (default: 0); it effectively limits the maximum size of loops found. Setting it to 0 results in an unlimited search depth.

The $-kso$ option generates a list of base pairs that link the loops in space, forming so-called kissing loops (see S.XL). This output analyzes all base pairs, including those that form pseudoknots (even if they are removed from the structure). By default, only canonical base pairs that form kissing loops are searched, but you can change this with the $-nock$ option to include non-canonical base pairs. The output file contains information about the secondary structures of the base pairs that form kissing loops.

The $-ldo$ option produces information about the structural units of secondary motifs called “ladders” (see S.XXXV) and “pseudoknots” (see S.XLI). The output file will contain information about the nucleotides included in the ladders/pseudoknots, the number of hydrogen bonds, their length, range, and gain. Depending on how you define a ladder, ladders are formed either exclusively from canonical base pairs or from base pairs of any type. By default, DSSNA forms ladders only from canonical base pairs, but you can override this behavior with the $-nocl$ option, forcing DSSNA to form ladders from base pairs of any type (but not pseudo-pairs).

If you want to analyze a pseudoknot-free structure, you should keep in mind that there is no universal method for detecting pseudoknots. DSSNA implements several pseudoknot detection algorithms, which are controlled by the $-knotmode$ parameter. By default, the $-knotmode$ *gromacs* mode is used, which is a modified optimization approach from Smit *et al.* (2008)[13]. Other available options represent other heuristic criteria from the same article (and follow the same logic): $-knotmode$ *ec* (elimination, conflicts), $-knotmode$ *eg* (elimination, gain), $-knotmode$ *io* (incremental, order), $-knotmode$ *il* (incremental, length), and $-knotmode$ *ir* (incremental, range). If you specify the $-knotmode$ *none* parameter, pseudoknots will not be searched for. Please note that the process of detecting pseudoknots and the process of eliminating them from the structure are two different processes.

After removing pseudoknots from the structure (if required), the algorithm by default attempts to resaturate ladders. Resaturation is the process of re-examining ladder structures and pseudoknots after their final identification, attempting to identify pseudoknots that do not form knots with ladders and restore them to ladder status. The name "resaturation" itself reflects the fact that without it, the structure could be undersaturated, potentially revealing too many pseudoknots. You can disable resaturation with the $-nores$ option.

Typically, immediately after pseudoknots are identified, the base pairs that form the pseudoknots are removed from the analysis. The full list of base pairs, with and without pseudoknots, is used only when searching for kissing loops and minor motifs (their identification requires full lists of base pairs). If you need all base pairs for your analysis, even those that form pseudoknots, you can use the $-nofix$ option to prevent the program from removing pseudoknot base pairs.

The $-dbno$ option generates dot-bracket notation (see S.XLII), a method for writing loops, ladders, and pseudoknots using dots, brackets, and letters of the English alphabet. Dot-bracket notation uses a forced single-letter notation for nucleotides. The dot-bracket notation generated by DSSNA can be subsequently visualized using various tools such as VARNA[15] or its Python interface varnaapi[15], facilitating the analysis of time-dependent changes in nucleic acid secondary structure. By default, pseudoknots are included (if detected) in the dot-bracket notation, but if you do not need them, use the $-nospn$ option to hide pseudoknots from the output file.

The $-sso$ option generates an output file containing information about the sequential chain of nucleotides that do not form loops, helices, and/or stems. Such sequences are called single-stranded fragments (see S.XXXII). The output file will contain information about the size of such a fragment and the nucleotides contained in each fragment.

The $-mto$ option outputs information about multiplets (see S.XXXIII), their sizes, and the nucleotides they comprise. The $-md$ option sets the maximum multiplet search depth, effectively limiting their size (useful if you are only interested in multiplets up to a fixed size). A value of 0 allows unlimited search depth.

The $-bbo$ option generates information related to the sugar-phosphate backbone geometry (see S.XLV). The output file specifies the nucleotides for each frame, whether a turn or break follows that nucleotide, and indicates torsional and virtual angles. You can choose between two sugar type detection modes (see S.L) using the $-sugarmode$ *x*3*dna* option (for sugar type detection as in X3DNA) and the $-sugarmode$ *curves* option (for sugar type detection as in Curves+).

The $-sao$ option analyzes and outputs splayed apart nucleotide conformation (see S.XLIV). The output file will contain data on the pair of splayed apart nucleotides, the angle and length between them, and the ratio of the normal to the vector based on the O3' and PO1'/PO2' oxygen positions. The $-saang$ flag controls the angle cutoff in the splayed apart conformation selection criterion.

The $-mmo$ option generates information on minor motifs (see S.XXXIV). The output file contains the name of the detected minor motifs, along with information on the nucleotides that form them. Minor motifs in DSSNA are written in the format "<interacting nucleotide> | <base pair>". By default, DSSNA considers the frequently occurring A-minor motifs, which often stabilize the tertiary structure of RNA. However, you can use the $-mn$ option to specify single-letter nucleotide designations relative to which minor motifs will be searched (select $-mn$ $N$ if you want to select all possible nucleotide types).

The $-gto$ option produces a file containing data on the G-tetrads contained in a nucleic acid structure (see S.XXXVI). Functionally, this option is equivalent to the option that generates multiplets, but only those that can be identified as G-tetrads. Algorithmically, DSSNA attempts to extract subsets of G-tetrads from the set of generated multiplets.

The $-imo$ option finds i-motifs in a nucleic acid structure (see S.XLIII). The output file will list the unique identifier of each detected i-motif and the nucleotides involved in its formation. DSSNA searches for i-motif helices and, based on the available data, analyzes the helix to extract strands and loops from it.

The $-rzo$ option generates information related to a special tertiary structure — the ribose zipper (see S.LII). The output file will list the contiguous nucleotide sequences that form ribose zippers (as well as their total size). DSSNA has two zipper search modes, each differing in how a zipper bond is defined: the $-zipmode\ default$ mode defines a zipper bond as the presence of a strict hydrogen bond between the O2' atoms of a base pair, while the simplified $-zipmode\ distance$ mode only requires that the distance between the O2' atoms of a base pair does not exceed the hydrogen bond threshold.

For a quick summary of all available options, please refer to Table S8. To view all standard default values that can be changed by the user, please refer to Table S9.

Table S8: A brief summary of the options that generate the output file in DSSNA.

| **Option name** | **Function** |
|---|---|
| -o | Nucleotide frames |
| -vo | Nucleotide virtual bonds |
| -hbo | Hydrogen bonds |
| -bpfo | Base pairs (location) |
| -bpro | Base pairs (rigid frames) |
| -bpso | Base pairs (secondary structures) |
| -bpvo | Base pairs (virtual bonds) |
| -bpoo | Base pairs (occupancy) |
| -bpmo | Base pairs (most stable lifetimes) |
| -dso | Dinucleotide and helical steps |
| -ho | Helices |
| -ldo | Ladders and pseudoknots |
| -dbno | Dot-bracket notation |
| -bso | Base-stacking interactions |
| -bsoo | Base-stacking interactions (occupancy) |
| -bsmo | Base-stacking interactions (most stable lifetimes) |
| -bco | Atom-base capping interactions |
| -lo | Loops |
| -sso | Single-stranded fragments |
| -mto | Multiplets |
| -bbo | Backbone |
| -sao | Splayed apart nucleotides |
| -kso | Kissing loops |
| -mmo | Minor motifs |
| -gto | G-Tetrads |
| -rzo | Ribose zippers |
| -imo | I-motifs |

Table S9: Default option parameters in DSSNA.

| Option name | Description | Default parameter value |
|---|---|---|
| -energy | Acceptor partial-charge cutoff | $-0.05\ e$ |
| -hbdist | Hydrogen bond distance cutoff | 0.35 nm |
| -hbang | Hydrogen bond angle cutoff | 30° |
| -bpang | Base pair geometry assignment angle cutoff | 40° |
| -bcdist | Atom-base capping distance cutoff | 0.55 nm |
| -bcvs | Atom-base capping vertical separation cutoff | 0.35 nm |
| -bcang | Atom-base capping angle cutoff | 25° |
| -bsarea | Base-stacking overlap ratio cutoff | 0.1 (10%) |
| -bsdist | Base-stacking distance cutoff | 0.45 nm |
| -bsang1 | Base-stacking angle cutoff | 23° |
| -bsang2 | Base-stacking angle cutoff | 40° |
| -saang | Splayed apart angle cutoff | 85° |
| -ld | Maximum loop search depth | 0 (unlimited) |
| -md | Maximum multiplet search depth | 0 (unlimited) |
| -ba | Atom-base capping elements | O |
| -mn | Nucleotide types for minor motif analysis | A |
| -sugarmode | Sugar type analysis mode | x3dna |
| -knotmode | Pseudoknot detection mode | gromacs |
| -bssmode | Base-stacking search mode | overlap |
| -zipmode | Ribose zipper search mode | default |
| -mapmode | Format for occupancy/most stable lifetime output | relation |
| -bpmmode | Base pair types for occupancy/lifetime analysis | canon |

### Molecular dynamics

Molecular dynamics (MD) modeling was performed with the software package GROMACS 2026[4]; the amber14sb+OL3 force field[21–23] was used for RNA, and TIP3P[24] was used as the water model. The resulting system was placed in a periodic water box so that at least 25 Å separated the RNA from the box walls. The resulting box was then minimized using the steepest descent algorithm. The system obtained as a result of minimization was subjected to charge neutralization by adding 150 mM NaCl, bringing the total charge of the system to zero. The neutralized system was again subjected to energy minimization using the steepest descent algorithm. Next, the system was equilibrated using a two-stage approach. At the first stage, all heavy atoms of RNA were restrained to their initial positions using an additional energy term (posres), and at the start of equilibration the temperature (particle velocity distribution) was taken from the Maxwell–Boltzmann distribution at 310 K. The system was equilibrated for 5 ns at this temperature. The integration step was 2 fs; the V-rescale thermostat[25] (coupling time 1 ps and coupling groups RNA and Water_and_ions) and the C-rescale barostat[26] (coupling time 1 ps) were used. At the second stage, the additional restraining potential was removed, and all components of the system could move freely. During this stage, the Nosé-Hoover thermostat[27] (coupling time 2 ps and coupling groups RNA and Water_and_ions) and the Parrinello-Rahman barostat[28] (coupling time 4 ps) were used, and the system was equilibrated for 10 ns. The final state obtained as a result of the two-stage equilibration was used as the starting point for the production dynamics. Molecular dynamics was carried out for 1000 ns using the same set of parameters as for the second stage of equilibration.

### Additional graphs and figures

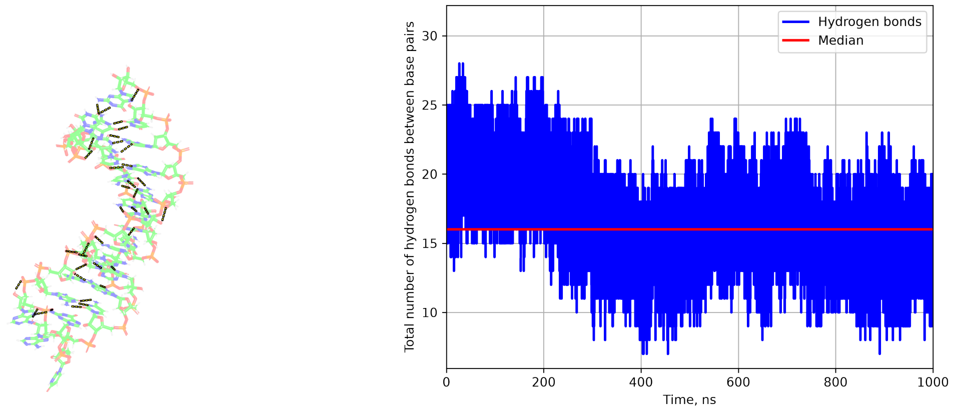


(a) (b)

Figure S22: (S22a) Visualization of the hydrogen bond network within the GUAA tetraloop mutant 1MSY. (S22b) Graph of the total number of hydrogen bonds between base pairs versus time in the trajectory of the GUAA tetraloop mutant 1MSY.

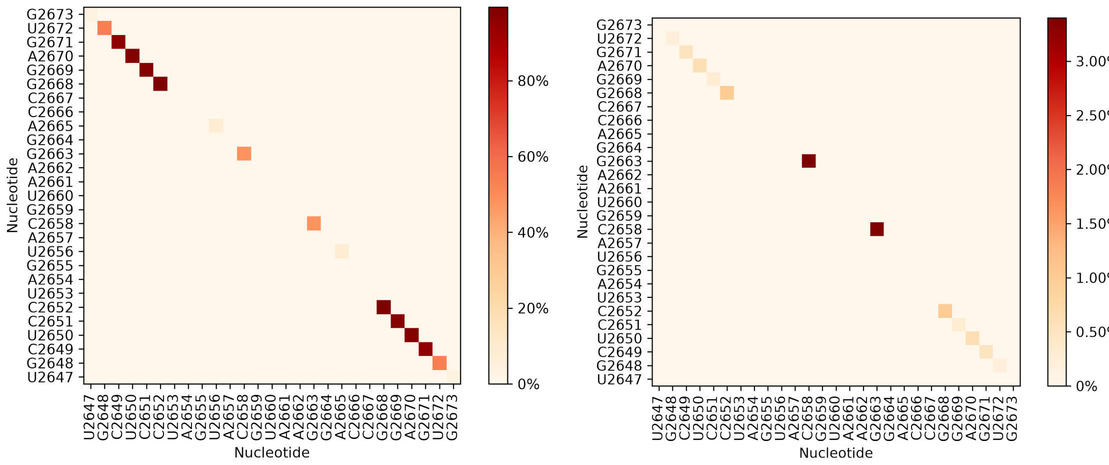


(a) (b)

Figure S23: (S23a) Heat map of base pair occupancies and (S23b) most stable lifetimes in the trajectory of the GUAA tetraloop mutant 1MSY. The values represent the normalized ratio of the lifetime to the total trajectory duration.

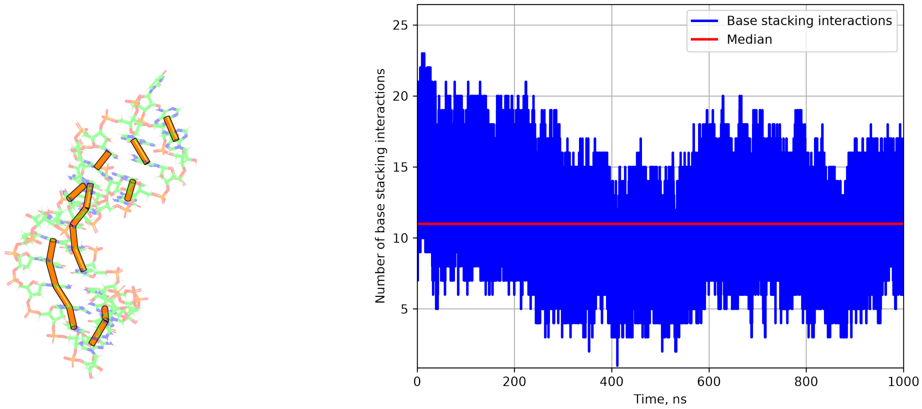


(a) (b)

Figure S24: (S24a) Visualization of base-stacking interactions within the GUAA tetraloop mutant 1MSY. (S24b) Graph of the number of base-stacking interactions versus time in the trajectory of the GUAA tetraloop mutant 1MSY.

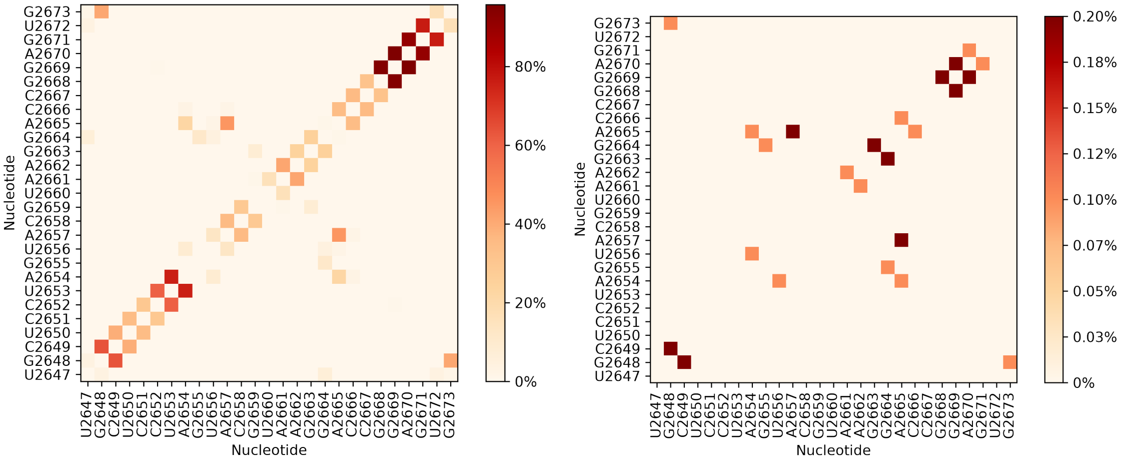


(a) (b)

Figure S25: (S25a) Heat map of base-stacking occupancies and (S25b) most stable lifetimes in the trajectory of the GUAA tetraloop mutant 1MSY. The values represent the normalized ratio of the lifetime to the total trajectory duration.

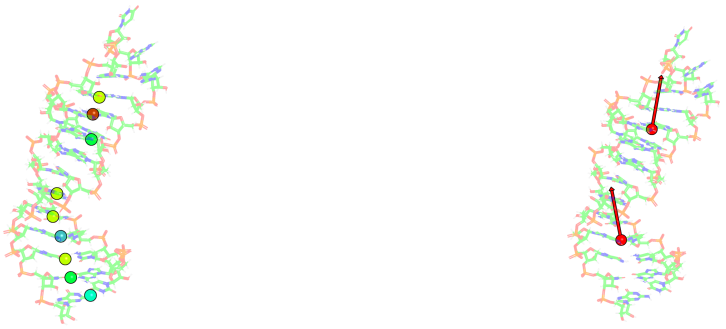

(a) (b)

Figure S26: (S26a) Visualization of base pair origins within the GUAA tetraloop mutant 1MSY. (S26b) Visualization of helical vectors within the GUAA tetraloop mutant 1MSY.
